# Conserved metabolic adaptation mechanisms in cancer cells and yeast against mitochondrial dysfunction indicate an additional role of aerobic glycolysis for cell survival

**DOI:** 10.1101/2025.02.16.637991

**Authors:** Akane Sawai, Takeo Taniguchi, Kohsuke Noguchi, Taisuke Seike, Nobuyuki Okahashi, Masak Takaine, Fumio Matsuda

**Author notes:** **Correspondence:** Fumio Matsuda, Department of Bioinformatic Engineering, Graduate School of Information Science and Technology, Osaka University, 1-5 Yamadaoka, Suita, Osaka 565-0871, Japan.

## Abstract

Eukaryotic cells primarily generate ATP through oxidative phosphorylation and substrate-level phosphorylation. Despite the superior efficiency of oxidative phosphorylation, eukaryotic cells often utilize both pathways as aerobic glycolysis even in the presence of oxygen. However, its role in cell survival remains poorly understood. In this study, aerobic glycolysis of the Warburg effect in breast cancer cells (MCF7) and the Crabtree effect in a laboratory strain of *Saccharomyces cerevisiae* (S288C) were compared following treatment with electron transport chain inhibitors, including FCCP, antimycin A, and oligomycin. MCF7 and S288C exhibited strikingly similar metabolic rewiring toward substrate-level phosphorylation against the inhibitor treatment, suggesting that mitochondrial oxidative phosphorylation and cytosolic substrate-level phosphorylation communicate through a common mechanism. Measurements of mitochondrial membrane potential (MMP) and ATP concentration further indicated that cytosolic ATP was transported into the mitochondria under conditions of reduced electron transport chain activity. This ATP was likely utilized by the reverse mode of H^+^/ATPase to maintain the MMP which contributed to avoiding programmed cell death. These results suggest that the ATP supply to mitochondria plays a conserved role in aerobic glycolysis across yeast and mammalian cancer cells. This mechanism likely contributes to cell survival under conditions of fluctuating oxygen availability.

## Introduction

Eukaryotic cells primarily produce ATP through two pathways: oxidative phosphorylation in the mitochondria and substrate-level phosphorylation in the cytosol [1]. Oxidative phosphorylation efficiently generates 32–36 ATP molecules through complete oxidation of one glucose molecule. In contrast, substrate-level phosphorylation produces only two ATP molecules during the conversion of glucose to lactate or ethanol, without requiring oxygen. Despite the high efficiency of oxidative phosphorylation, eukaryotic cells often utilize both pathways simultaneously, even in the presence of oxygen. For instance, cancer cells preferentially rely on substrate-level phosphorylation, a phenomenon known as the Warburg effect, is considered a hallmark of cancer [2, 3]. However, oxidative phosphorylation remains active in many cultured cancer cells, with some supplying more than 50% of their ATP through oxidative phosphorylation under aerobic condition [4–6]. Similarly, Crabtree-positive yeasts, such as *Saccharomyces cerevisiae*, have evolved the ability to perform substrate-level phosphorylation even under aerobic conditions [7].

The Warburg effect in human cancer cells and the Crabtree effect in *S. cerevisiae* represent two manifestations of aerobic glycolysis [1, 8]. A question regarding aerobic glycolysis is why cancer and *S. cerevisiae* deliberately rely on the inefficient process of substrate-level phosphorylation, even under aerobic conditions. Several roles for aerobic glycolysis have been proposed, such as rapid ATP and substrate supply to support cell proliferation [9], modulation of the cancer microenvironment through lactate excretion [10], and a possible advantage related to the prevalence from an upper limit on Gibbs energy dissipation [11]. However, further investigation is needed to understand the multifaceted role of aerobic glycolysis.

Another question regarding aerobic glycolysis is whether the metabolic regulatory mechanisms are conserved between mammalian cancer cells and unicellular *S. cerevisiae*. Previous studies have demonstrated that complex regulatory mechanisms govern the interaction between oxidative phosphorylation and substrate-level phosphorylation, as perturbations in one pathway can lead to compensatory changes in the other [12, 13]. For instance, in melanoma cells, a low-dose treatment with complex I inhibitors activates substrate-level phosphorylation [14]. Similarly, in *S. cerevisiae*, increased NADH oxidation has been shown to reduce the overflow metabolism [15]. Additionally, the overexpression of Hap4, a positive transcriptional regulator of respiratory metabolism genes, modulates the flux distribution between respiration and fermentation in *S. cerevisiae* [16]. If the regulatory mechanisms driving aerobic glycolysis are shared between human cancer and *S. cerevisiae*, this suggests that the role of aerobic glycolysis may be more intrinsic to metabolic function and essential for the survival of eukaryotic cells.

This study investigated the physiological significance of aerobic glycolysis and the interaction mechanisms between cytoplasm and mitochondria during aerobic glycolysis. We employed two complementary approaches to address these research objectives. First, we conducted a comparative analysis of aerobic glycolysis between human breast cancer cells (MCF7) and the *S. cerevisiae* laboratory strain (S288C). MCF7 cells were selected due to their characteristic ATP production profile, with approximately 30% of ATP generated through oxidative phosphorylation under aerobic conditions [17]. Second, we examined the cytoplasm-mitochondria interactions during small perturbations induced by low doses of three electron transport chain inhibitors: FCCP, antimycin A, and oligomycin. The results demonstrated remarkably conserved metabolic responses to oxidative phosphorylation dysfunction between cancer cells and yeast at the flux, metabolome, and gene expression levels. Furthermore, our data indicated that upon suppression of electron transport chain activity, cytosolic ATP was transported to the mitochondria and utilized by the reverse mode of H^+^/ATPase to maintain mitochondrial membrane potential (MMP). Given that decreased MMP promotes programmed cell death, this mechanism likely plays a crucial role in cell survival under conditions of variable oxygen availability.

## Results

### Similar metabolic rewiring against respiratory chain inhibitor treatment between breast cancer cells, MCF7, and *S. cerevisiae* cells, S288C

To evaluate the effect of the respiratory chain inhibitor on cell proliferation, MCF7 cells, a human breast cancer cell line, were seeded in a dish containing passage medium and cultured for 15 h. Once the cells adhered to the dish, the medium was replaced with a metabolic analysis medium supplemented with specific respiratory chain inhibitors. This study utilized FCCP, antimycin A, and oligomycin to disrupt oxidative phosphorylation-dependent ATP synthesis (Fig. 1a): FCCP functions as an uncoupler by transporting protons across the mitochondrial inner membrane, dissipating the proton gradient. Antimycin A inhibits electron transport from cytochrome b to the cytochrome complex III. Oligomycin inhibits mitochondrial H^+^/ATPase, blocking both ATP synthesis and hydrolysis. The concentrations of FCCP, antimycin A, and oligomycin used in this study were similar levels to the standard protocol for the FluxAnalyzer (FCCP: 0.125–2.0 µM, antimycin A 500 nM, and oligomycin 1500 nM) [18].

**Figure 1.**
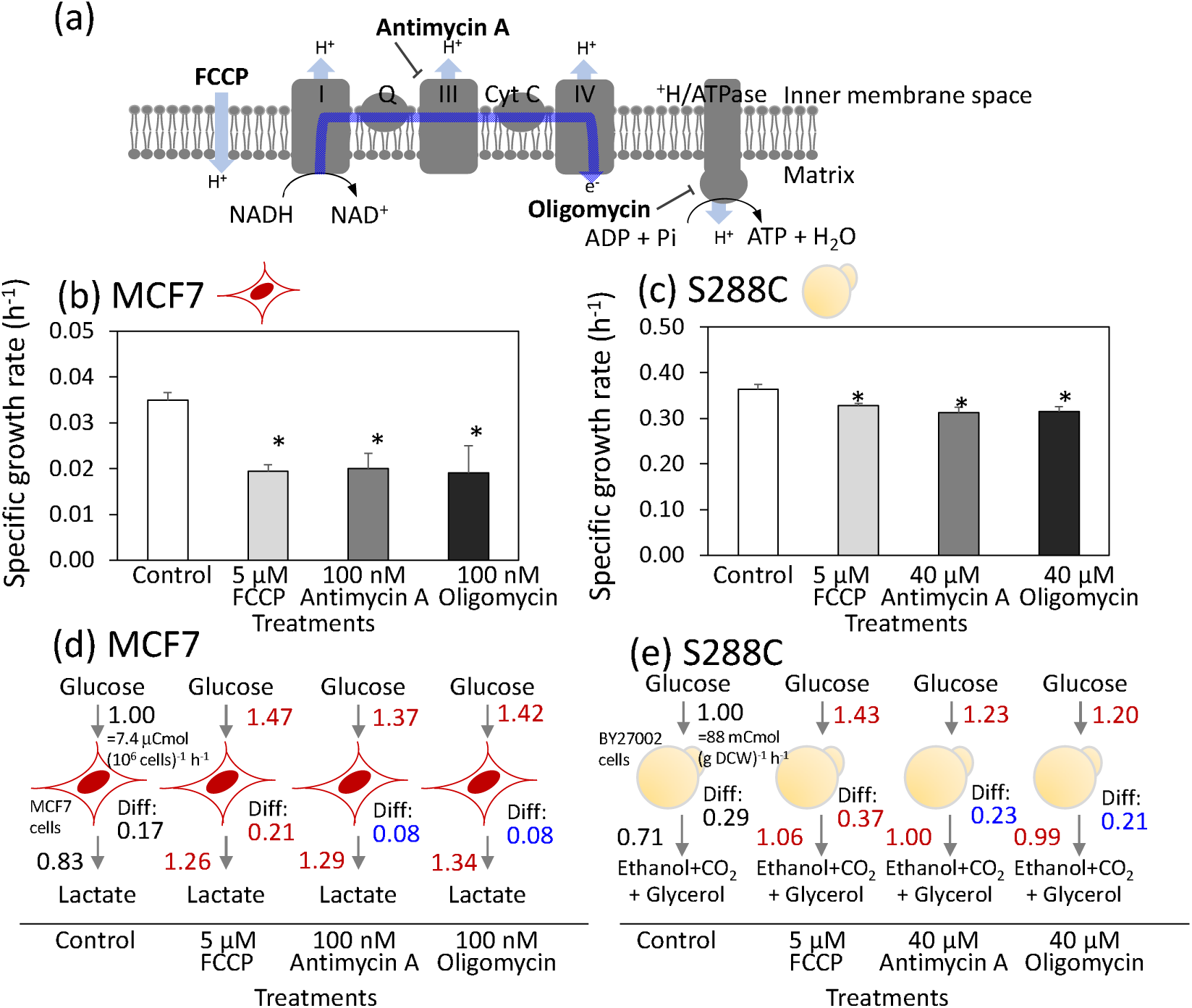
Similar metabolic rewiring against respiratory chain inhibitor treatment between human breast cancer cells, MCF7, and *Saccharomyces cerevisiae,* S288C. (a) Targets of respiratory chain inhibitors in human mitochondria (b, c) Effect of FCCP, antimycin A, and oligomycin treatment on specific growth rates of (b) MCF7 and (c) S288C. All results were obtained from triplicate cultures and described as mean ± SD. Asterisks indicate *p*-value < 0.05 by two-sided *t*-test. (d, e) Metabolic rewiring of (d) MCF7 and (e) S288C against the inhibitor treatment. Carbon balances were calculated from the specific rates for glucose consumption and product excretion. All values represent the carbon flux level relative to the carbon uptake level of the control culture. Differences in carbon balance (Diff) between glucose uptake and product excretion are also shown. All results were obtained from triplicate cultures and described as mean values. Red and blue letters indicate a significant increase and decrease with *p*-value < 0.05 by two-sided *t*-test compared to the control.

The specific growth rate of the control MCF7 culture during the log phase was 0.035±0.002 h□¹ (Fig. 1b). When treated with 5 μM FCCP, 100 nM antimycin A, and 100 nM oligomycin, the specific growth rate decreased to approximately half that of the control culture (Fig. 1b). These results suggest that the respiratory chain was partially inhibited but not completely stopped by these inhibitor concentrations.

For comparison, cells of a laboratory strain of *S. cerevisiae* (S288C) were batch-cultured under aerobic conditions in 6-well plates. Inhibitors were added 4 h after the start of cultivation. The specific growth rate of the control culture was determined to be 0.37±0.01 h□¹ from the growth curve (Fig. 1c and Table S1). S288C displayed greater tolerance to the inhibitor treatment, as 5 μM of FCCP reduced growth by only 9.7% (Fig. 1c). Similarly, growth inhibitions of 19 and 22% were observed for antimycin A and oligomycin, respectively, even at the highest tested concentration of 40 μM (Fig. 1c). As partial growth inhibition was achieved in both MCF7 and S228C cells, these inhibitor concentrations were used consistently throughout the study.

To investigate the metabolic rewiring induced by inhibitor treatment, MCF7 cells were cultured under identical conditions. The concentrations of glucose and lactate in the medium were measured over time to calculate the mass balance during the log phase (Fig. 1d and Table S2). The specific rates for glucose uptake and lactate production in the control culture were determined to be 1.23±0.03 and 2.05±0.02 μmol (10□ cells)□¹ h□¹, respectively. These rates corresponded to the carbon uptake and production rates of 7.40±0.12 and 6.15±0.06 μCmol (10□ cells)□¹ h□¹, as glucose and lactate contain six and three carbon atoms per molecule, respectively. The relative number of carbon atoms excreted as lactate was 0.83 when the carbon uptake rate was set to 1.00. The active production of lactate from glucose even under aerobic conditions is referred to as aerobic glycolysis, also known as the Warburg effect. FCCP treatment significantly enhanced both carbon uptake and lactate excretion, with relative levels of 1.47 and 1.26, respectively, compared to the control (Fig. 1d). These findings indicate that FCCP treatment induces metabolic rewiring, characterized by enhanced aerobic glycolysis. Similar increases in aerobic glycolysis were observed upon treatment with antimycin A and oligomycin (Fig. 1d).

The difference in the carbon balance (Diff) between glucose uptake and lactate excretion was further examined. For control MCF7 cells, the Diff value was calculated to be 0.17 (Fig. 1d). Most of the carbon atoms were consumed for oxidative phosphorylation via the TCA cycle in the mitochondria, as well as for the synthesis of cell components such as amino acids and lipids. Analysis of the inhibitor-treated cells showed that the Diff value increased to 0.21 with FCCP treatment and decreased to 0.08 with antimycin A and oligomycin treatment (Fig. 1d). The reduction of the Diff value for antimycin A- and oligomycin-treated cells reflects decreased carbon consumption for both the TCA cycle and biosynthesis of cell components, due to inhibition of the respiratory chain. In contrast, the elevated Diff levels in FCCP-treated cells suggest an increased flux through the TCA cycle, as well as in the respiratory chain. Since FCCP is an uncoupler of the proton gradient in the mitochondria, this activation of the respiratory chain likely serves to maintain the MMP.

A similar analysis was performed for S288C (Fig. 1e and Table S1). The carbon uptake rate for glucose in the control was calculated to be 88 mCmol g DCW□¹ h□¹. Among these, the carbon consumption attributed to ethanol, CO_2_, and glycerol production accounted for 0.71 (Fig. 1e). This active aerobic glycolysis in S288C is also known as the Crabtree effect. Moreover, the metabolic rewiring in S288C following inhibitor treatment was nearly identical to that observed in MCF7 cells (Fig. 1e).

### The control mechanisms governing metabolic flux are shared between human breast cancer cells, MCF7, and *S. cerevisiae*, S288C

To estimate the metabolic control mechanisms responsible for metabolic rewiring, a metabolic profiling analysis was conducted. MCF7 and S288C cells were cultivated under the aforementioned conditions. Cells were collected 3 and 2 h after inhibitor treatment, respectively, and preserved for targeted metabolome analysis using LC-MS/MS and GC-MS. A metabolic profile dataset, including 23 metabolites involved in central carbon metabolism, was used for the analysis (Table S3). Figure 2a shows the amounts of metabolites relative to the control samples.

**Figure 2.**
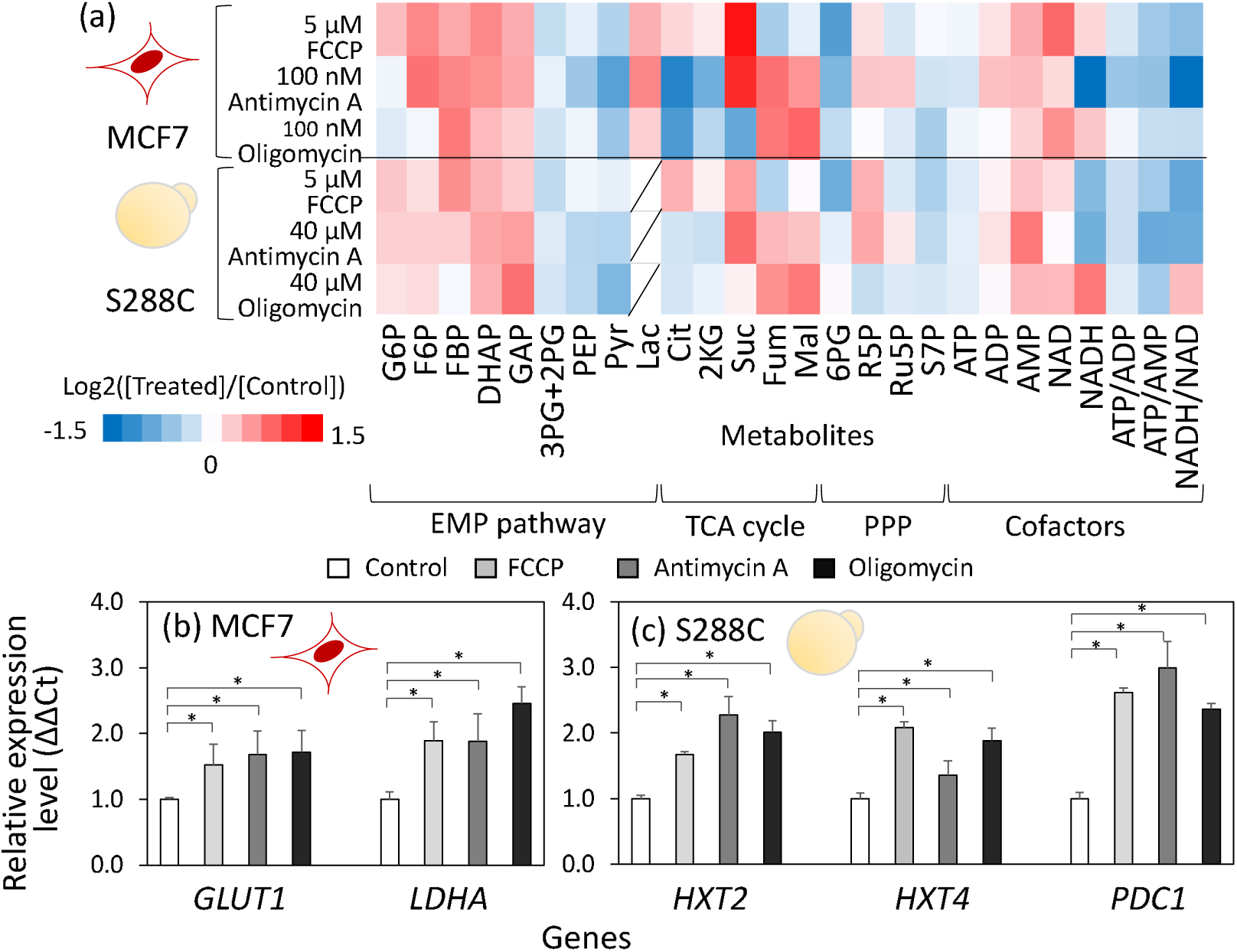
Similar metabolic responses against respiratory chain inhibitor treatment between human breast cancer cells, MCF7, and *S. cerevisiae*, S288C. (a) Metabolic profiles in inhibitor-treated human breast cancer cells, MCF7, and *S. cerevisiae,* S288C. Intracellular metabolites in MCF7 and S288C cells were collected at 3 and 2 h after inhibitor treatment, respectively, and served for the targeted metabolome analysis. All results were obtained from duplicate or triplicate cultures. The relative amounts of metabolites to the control are shown in heatmap. (b,c) Effect of FCCP, antimycin A, and oligomycin treatment on gene expression of (b) MCF7 and (c) S288C cells. RNA was collected from MCF7 and S288C cells at 3 and 2 hours after inhibitor treatment, respectively, and served for the quantitative PCR analysis. The relative expression levels to the control were shown. All results were obtained from triplicate cultures and described as Mean±SD. Asterisks indicate *p*-value < 0.05 by two-sided t-test.

The metabolic profiles revealed that treatment with FCCP, antimycin A, and oligomycin commonly caused a reduction in the ATP content as well as the ATP/ADP and ATP/AMP ratios, in both MCF7 and S288C (Fig. 2a). This metabolic state indicated that the inhibition of the respiratory chain decreased ATP supply. Moreover, the inhibitor treatments commonly induced an increase in metabolites in the upper part of the Embden-Meyerhof-Parnas (EMP) pathway, including fructose-6-phosphate (F6P), fructose-1,6-bisphosphate (FBP), dihydroxyacetone phosphate (DHAP), and glyceraldehyde-3-phosphate (GAP), while a decrease in metabolites in the lower part of the EMP pathway, including 3-phosphoglycerate (3PG) + 2-phosphoglycerate (2PG), phosphoenolpyruvate (PEP), and pyruvate (Pyr). The metabolic profile is a signature of elevated flux level of the EMP pathway because similar metabolic responses were observed during glucose pulse experiments in *S. cerevisiae* [19]. In these experiments, under glucose-limited conditions, the addition of a high concentration of glucose suddenly elevated the metabolic flux in the EMP pathway, leading to an accumulation of upper EMP pathway metabolites, including FBP, which is an allosteric activator of pyruvate kinase that catalyzes the final step of the EMP pathway. The accumulation of FBP caused the activation of pyruvate kinase, promoting the elimination of intermediates in the lower part of the EMP pathway, including 2PG, 3PG, and PEP [20, 21].

In contrast to the common response in the EMP pathway, the metabolite profiles of the TCA cycle differed between the inhibitors. In MCF7 cells, FCCP treatment caused an elevation of citrate (Cit) and 2-ketoglutarate (2KG) levels, while the levels of fumarate (Fum) and malate (Mal) decreased (Fig. 2a). Conversely, in cells treated with antimycin A and oligomycin, a decrease in Cit and 2KG and an increase in Fum and Mal were observed (Fig.2a). A similar trend was observed in S288C (Fig. 2a). These metabolic responses suggested that the entry point of the TCA cycle was upregulated by FCCP treatment and downregulated by antimycin A and oligomycin treatments, which was consistent with the Diff levels in metabolic rewiring shown in Figure 1d and e.

Next, the control mechanisms of metabolic rewiring were investigated using gene expression analyses. Quantitative PCR analysis of MCF7 cells, conducted 3 h after inhibitor treatment, revealed that the expression of *GLUT1*, which is reported to be a highly expressed glucose transporter gene in cancer cell lines [22], increased significantly by 1.5-to 1.7-fold following inhibitor treatment (Fig. 2b). Moreover, the expression of *LDHA*, a major isozyme of lactate dehydrogenase (LDH), increased 1.8 to 2.4-folds after inhibitor treatment (Fig. 2b). Gene expression analysis of S288C also showed that inhibitor treatment commonly upregulated the expression levels of the high-affinity glucose transporters *HXT2* and *HXT4* [23] and *PDC1*, which encodes the major isoenzyme of pyruvate decarboxylase (PDC), involved in the biosynthesis of ethanol from pyruvate (Fig. 2c).

These results were consistent with the metabolome data. The low ATP/AMP ratio in the inhibitor-treated cells likely activated AMP-activated protein kinase (AMPK) which is a positive regulator of *GLUT1* and *LDHA* expression [24]. Moreover, these findings suggest that the control mechanisms governing metabolic flux in the EMP pathway and the TCA cycle are shared between human breast cancer cells, MCF7, and *S. cerevisiae*, S288C, as evidenced by the similar changes in metabolic flux, metabolite accumulation, and gene expression levels.

### Constant mitochondrial membrane potential against respiratory chain inhibitor treatment

A decrease in MMP is closely related to cell survival through the induction of apoptosis [25]. Since the inhibition of the respiratory chain is expected to affect the MMP, we investigated the effect of respiratory chain inhibitor treatment on the MMP. To analyze MCF7 and S288C, cultures were incubated for 3 and 2 h, respectively, before being treated with MitoTracker reagent for fluorescence imaging analysis (Fig. S1). Fluorescence imaging analysis showed no significant change in the mean fluorescence levels after inhibitor treatment in either MCF7 or S288C (Fig. S2, Fig. 3a and b). While the lack of effect on MMP by the respiratory chain inhibitor treatment was counterintuitive, it can be explained by the low doses of inhibitors and the existence of some mechanisms to maintain MMP. These results suggest that the potential purpose of the metabolic rewiring was to maintain the MMP at a constant level despite inhibitor treatment.

**Figure 3.**
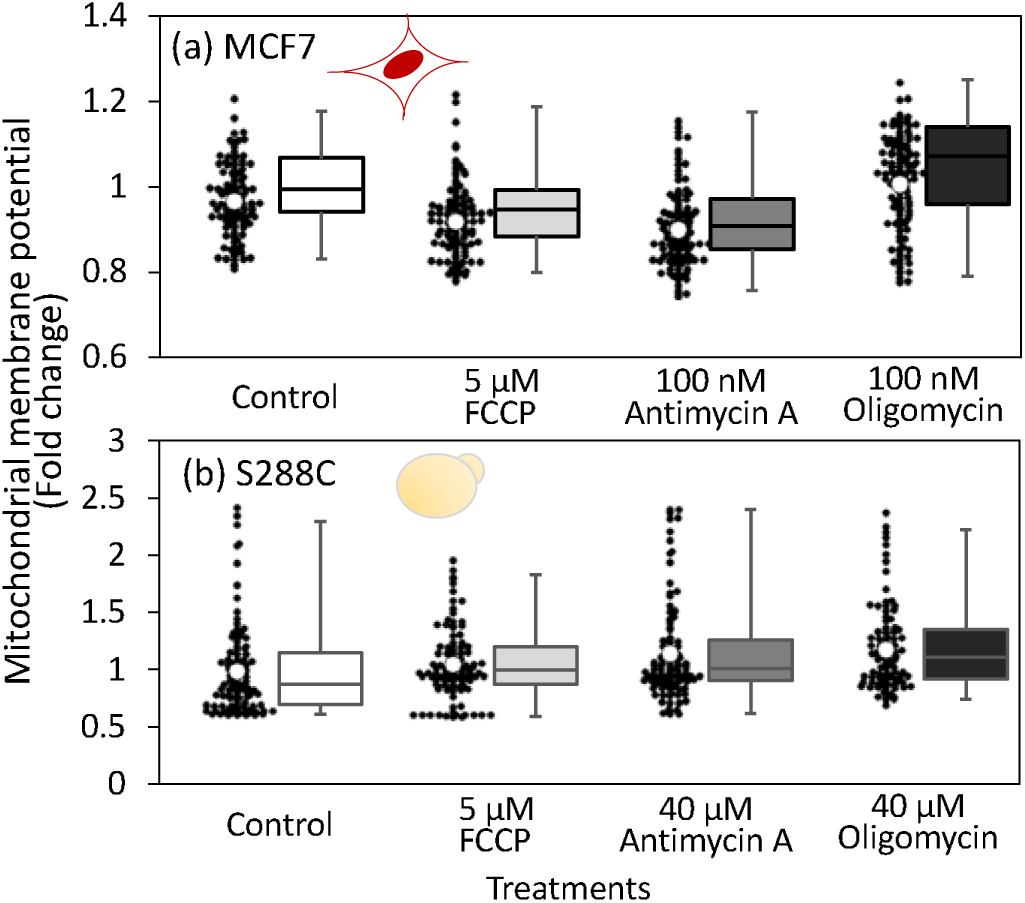
Effect of FCCP, antimycin A, and oligomycin treatment on mitochondrial membrane potential levels. MitoTracker reagent was applied to MCF7 and S288C cells 3 and 2 h after inhibitor treatment, respectively, followed by fluorescent microscopy analysis. Black dots indicate the mean fluorescence intensity of each individual cell, calculated from the fluorescence images. White circles represent the mean intensity levels of all investigated cells relative to the control. A box plot is also used to illustrate the distribution intensity values. No significant change in MMP was observed, as determined by a two-sided *t*-test with α = 0.05.

### Activation of mitochondrial ATP consumption by FCCP treatment

To investigate the mechanisms underlying the robust MMP, ATP levels in the cytoplasm and mitochondria of *S. cerevisiae* were measured using the ATP biosensor QUEEN-2m [26]. For this purpose, two *S. cerevisiae* strains expressing QUEEN-2m in the cytoplasm (YAS002) and mitochondria (YAS003) were generated from the BY18615 strain (Table S4 and Fig. S3). Both strains were cultured under the identical conditions. Cells were collected during the log phase and adhered to glass plates. Following the inhibitor treatment for 12 min, the fluorescence intensity of each cell was measured every 3 min for 120 min using a fluorescence microscope. ATP in the cytoplasm is denoted as ATP_Cyto_, and ATP in the mitochondria is denoted as ATP_Mito_.

Treatment with respiration inhibitors was expected to decrease mitochondrial ATP production. However, immediately after FCCP treatment, ATP_Mito_ levels increased by approximately 1.2 times, while ATP_Cyto_ levels in the cytoplasm decreased to approximately 0.8 times their initial value (Fig. 4a). This result suggests a downregulation of ATP transport from the mitochondria to the cytosol or activation of reverse ATP transport from the cytosol to the mitochondria. The latter response is plausible because FCCP is known to cause magnesium leakage from the mitochondria [27] and mitochondrial calcium accumulation [28], which triggers the activation of the ATP-Mg/Pi transporter and the accumulation of ATP in mitochondria. The accumulated ATP is thought to drive H^+^/ATPase in the reverse mode, generating a proton gradient as a proton pump [29]. The Pi generated by ATPase is expelled back into the cytoplasm by the ATP-Mg/Pi transporter.

**Figure 4.**
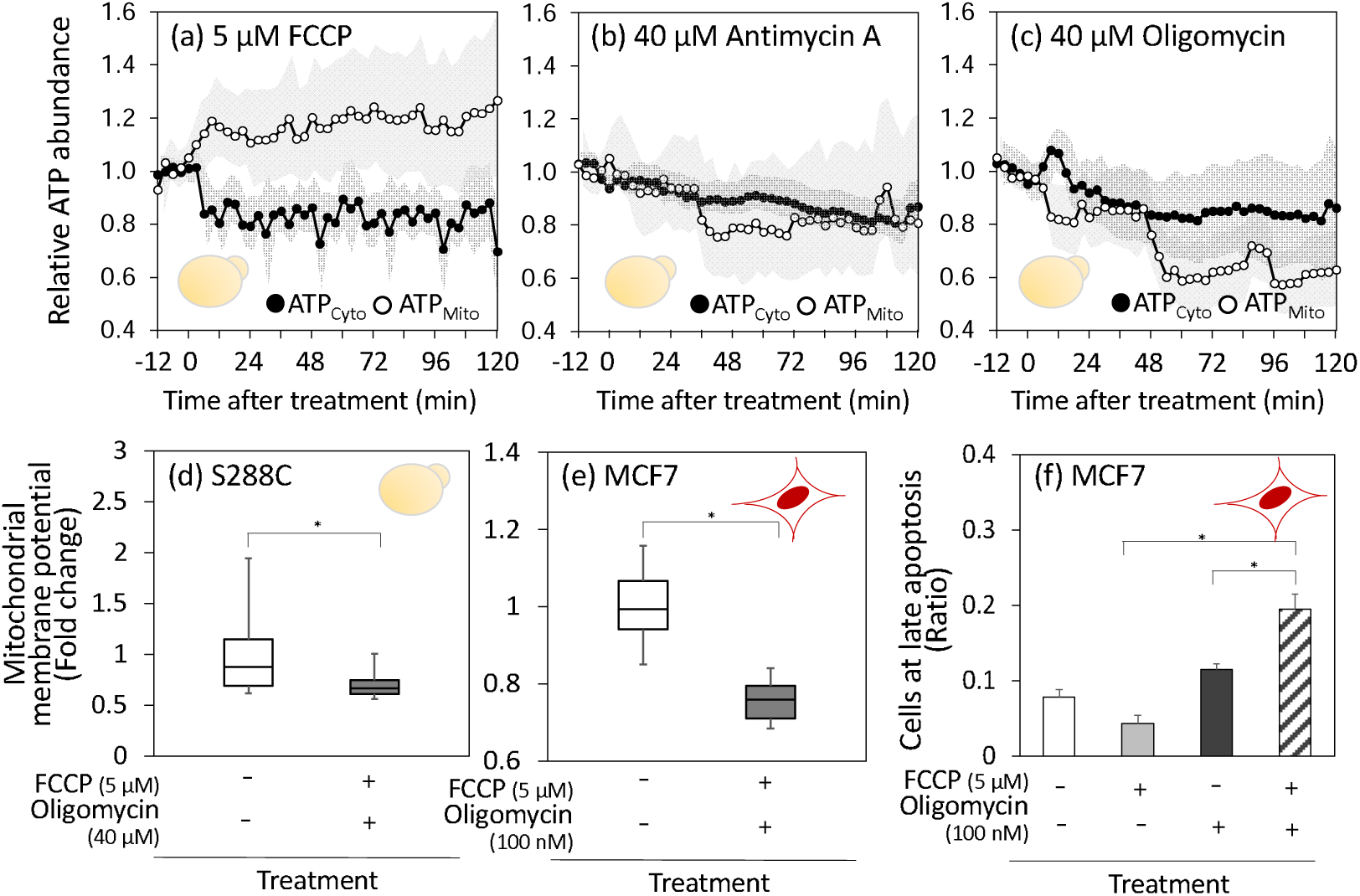
Mitochondrial ATP consumption by FCCP treatment via activation of reverse-mode H^+^/ATPase. (a–c) ATP abundances in the cytoplasm (ATP_Cyto_) and mitochondria (ATP_Mito_) of *S. cerevisiae* after (a) 5 µM FCCP, (b) 40 µM antimycin A, and (c) 40 µM oligomycin treatment. Cells from two *S. cerevisiae* strains expressing the ATP biosensor QUEEN-2m in the cytoplasm or mitochondria were collected during the log phase and adhered to glass plates. Following the inhibitor treatment for 12 min, the fluorescence intensity of each cell was measured every 3 min using fluorescence microscopy. For each cell, fluorescence intensity at the start of the experiment was set to 1.0. Mean values of more than 30 cells were shown as open and close circles. Gray shadow represents their standard deviations. (d–e) MMP after co-treatment with FCCP and oligomycin. (d) *S. cerevisiae* S288C and (e) human breast cancer cells (MCF7) were treated with inhibitors for 2 and 3 h, respectively, and analyzed using MitoTracker reagent. The box plot represent the distributions of more than 30 cells. (f) Induction of apoptosis by the inhibitor treatment. Human breast cancer cells MCF7 were treated with inhibitors for 24 h, and proportion of cells in late apoptosis were determined. Data was presented from triplicate cultures. Asterisks indicate *p*-value < 0.05 by two-sided *t*-test.

In the case of antimycin A treatment, there were no significant changes in ATP_Cyto_ or ATP_Mito_ levels (Fig. 4b). Subsequently, both ATP_Cyto_ and ATP_Mito_ levels gradually decreased in parallel, reaching 0.8 to 0.9 of their initial levels after 120 min. During oligomycin treatment, ATP_Mito_ levels briefly decreased and then dropped again at approximately 50 min to approximately half of the control levels (Fig. 4c). ATP_Cyto_ levels also gradually decreased. A possible interpretation of these behaviors is that mitochondrial ATP levels are maintained by reducing ATP transport from the mitochondria to the cytosol or by reverse ATP transport from the cytosol to the mitochondria.

To investigate the direction of ATP transport, the cells were co-treated with FCCP and oligomycin. Unlike the individual treatments shown in Figure 3, the co-treatment of FCCP and oligomycin reduced the MMP in S288C (Fig. 4d, Fig. S4). The result suggested that oligomycin inhibits ATP hydrolysis by H^+^/ATPase when treated simultaneously with FCCP, and that H^+^/ATPase activity contributes to the maintenance of the membrane potential. This decrease in MMP after co-treatment was also observed in MCF7 cells (Fig. 4e, Fig. S4). These results suggest that ATP transport from the cytoplasm to the mitochondria, coupled with H^+^/ATPase functioning in the reverse mode contributes to the robust MMP against inhibitor treatment.

Next, we investigated the relationship between the decreased MMP and programmed cell death. For MCF7, cells were collected 24 h after inhibitor treatment and subjected to apoptosis assays. The results showed that compared to individual FCCP or oligomycin treatments, co-treatment with FCCP and oligomycin increased the proportion of cells in the late stages of apoptosis by approximately 1.5-3.0 fold (Fig. 4f). This suggests that the decrease in MMP caused by co-treatment triggers cellular apoptosis. However, co-treatment failed to induce cell death in S288C (data not shown), probably because of the lower susceptibility of S288C to inhibitors (Fig. 1c).

### Mitochondrial membrane potential under hypoxic or low oxygen conditions

Under hypoxic conditions, the activity of the electron transport chain is reduced due to lack of oxygen as an electron acceptor, suggesting that H^+^/ATPase operates in the reverse mode to maintain the MMP [29–31]. To investigate the role of H^+^/ATPases, MCF7 and S288C cells were treated with oligomycin under hypoxic conditions. Unlike the normoxic conditions shown in Figure 3, the results in MCF7 cells showed that the MMP decreased by 27% with 100 nM oligomycin treatment in the hypoxic environment (Fig. 5a, Fig. S5).The result suggested that ATP hydrolysis by H^+^/ATPase was inhibited by oligomycin in the hypoxic condition. A similar decrease in the MMP was observed in oligomycin-treated S288C (Fig. 5b, Fig. S5).

**Figure 5.**
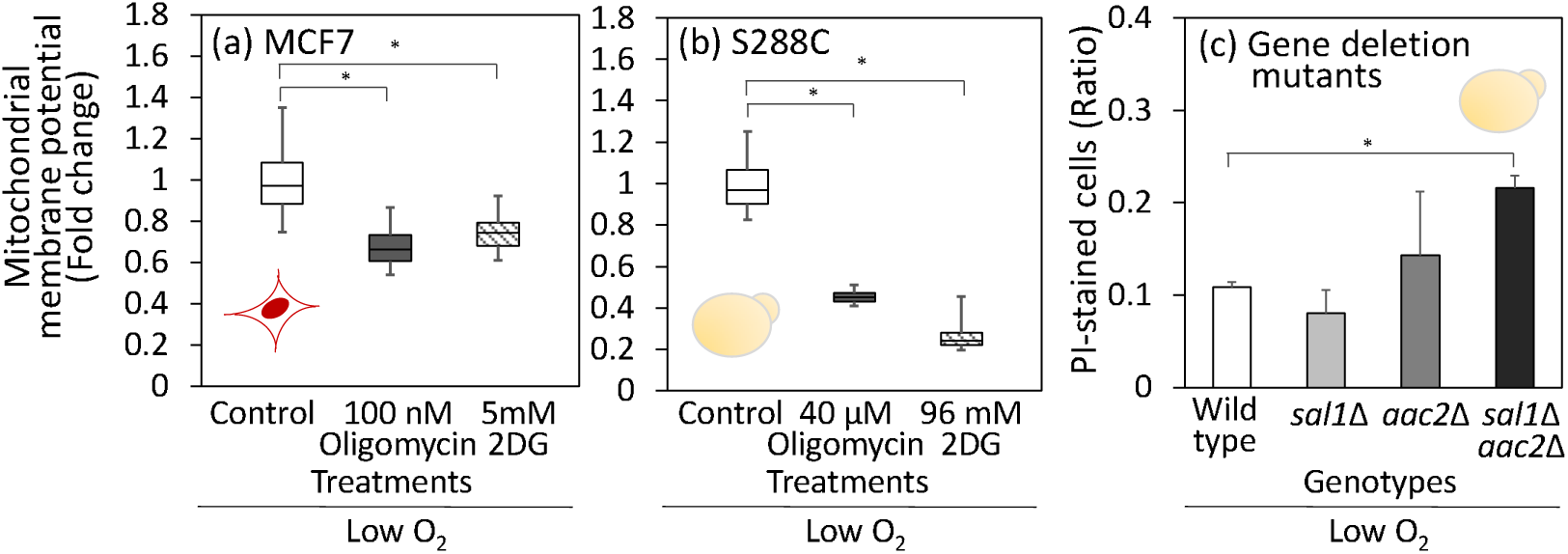
Role of reverse mode H^+^/ATPase under low oxygen condition. (a, b) Effect of oligomycin and 2-deoxyglucose (2DG) treatment on MMP levels. (a) MCF7 cells were cultured in the medium containing the inhibitors for 24 h at 1% O□, 5% CO□, 37 °C using a low-oxygen incubator. (b) For S288C, cells were cultured in the 1 mL of SD medium containing inhibitors and placed in an anaerobic chamber at 30 °C for 24 h. After treatment with MitoTracker reagent, fluorescence was observed while maintaining hypoxic conditions on a glass plate in the low-oxygen incubator. Box plots represent distribution of MMP for more than 30 cells. Data were obtained from triplicated cultures. (c) Effect of mitochondrial ATP carrier gene deletions on cell viability under the anaerobic condition. Wild type, *sal1*Δ, *aac2*Δ, and *sal1*Δ*aac2*Δ strains were cultured under the aforementioned anaerobic conditions for 24 h. Cell were stained with PI to examine dead cell rates through microscopic observation. Data were obtained from duplicated cultures. Asterisks indicate *p*-value < 0.05 by two-sided *t*-test.

Moreover, to assess the dependence on ATP transport from the cytoplasm to the mitochondria, 2-deoxyglucose (2DG) was used to inhibit the EMP pathway. The MMP of both MCF7 cells and S288C decreased significantly after 2DG treatment (Fig. 5a and b). These results support the hypothesis that the reverse mode of H^+^/ATPase and ATP derived from the substrate level phosphorylation contribute to maintaining MMP under low-oxygen conditions.

ADP/ATP carrier proteins in the inner mitochondrial membrane play a crucial role in ATP transport between the cytoplasm and mitochondria. Humans and *S. cerevisiae* commonly have three isoforms: *AAC1*, *AAC2*, and *AAC3* genes for the case of *S. cerevisiae.* It has been reported that Aac2 makes the largest contribution to ATP transport from the cytosol to the mitochondria [32]. Additionally, the calcium-dependent ATP-Mg/Pi carrier Sal1 contributes to the cytosol-to-mitochondrial ATP transport [33]. In this study, single and double gene deletion strains including *sal1*Δ, *aac2*Δ, and *sal1*Δ*aac2*Δ were constructed and cultured under the anaerobic conditions for 24 h. Cells were stained by propidium iodide (PI) to assess the dead cell rate (Fig. 5c). In the wild-type strain, the dead cell rate was 10.9%. Whereas the *sal1*Δ and *aac2*Δ strains showed no significant increase in dead cell rate, the *sal1*Δ*aac2*Δ double knockout strain exhibited significant increase, with a dead cell rate of 19.6% (Fig. 4c). These results suggest that ATP transport from the cytoplasm to the mitochondria plays an important role in maintaining cell viability of *S. cerevisiae* under anaerobic conditions.

## Discussion

Aerobic glycolysis is a metabolic state in which ATP regeneration occurs via both oxidative and substrate-level phosphorylation even in the presence of oxygen. Two well-known aerobic glycolysis are the Warburg effect in mammalian cancer cells and the Crabtree effect in the unicellular organism *S. cerevisiae* [1, 8]. This study showed that the metabolic responses to oxidative phosphorylation dysfunction caused by inhibitor treatment were remarkably similar between cancer cells and yeast at the flux, metabolome, and gene expression levels (Fig. 1 and 2). This similarity indicates that similar regulatory mechanisms are involved in the metabolic control of MCF7 and S288C cells. The major controlling points were the entry (glucose transporter) and exit (LDH and PDC) steps of the EMP pathway (Fig. 2b and c) as well as the entry steps of the TCA cycle (Fig. 2a). It has been reported that a low ATP/AMP ratio in inhibitor-treated cells upregulates the EMP pathway via AMPK [24]. This upregulation causes an accumulation of the EMP pathway intermediate FBP. The Ras-cAMP/PKA pathway appears to play a role in the subsequent adaptation, as both *S. cerevisiae* and human cancer cells share a conserved mechanism in which FBP binds to Cdc25/Sos1 and activates downstream Ras and cAMP/PKA pathways [34].

The striking similarity of the metabolic response also suggests that mitochondrial oxidative phosphorylation and cytoplasmic substrate-level phosphorylation are interconnected through a common mechanism in both mammalian cancer cells and unicellular yeast cells. The data presented in this study indicate that the ATP supply to mitochondria plays a crucial role in aerobic glycolysis. As its name implies, mitochondrial H^+^/ATPase can hydrolyze ATP and acts as a proton pump to generate membrane potential [29]. The reverse mode of H^+^/ATPase activity was inferred from the observation that the MMP remained unchanged in inhibitor-treated MCF7 and S288C cells (Fig. 3). Analysis using *S. cerevisiae* strains expressing the ATP sensor showed that cytoplasmic ATP levels decreased while mitochondrial ATP levels increased after FCCP treatment (Fig. 4a). Furthermore, simultaneous treatment with FCCP and oligomycin led to a decreased MMP in both *S. cerevisiae* S288C and human breast cancer cells MCF7 (Fig. 4d and e). Moreover, under hypoxic conditions, MMP decreased with either oligomycin treatment and the EMP pathway inhibition in both cell types (Fig. 5a and b).

Similar observations have been made in other mammalian cells. For example, in nerve cells, complete inhibition of complex I with rotenone still maintains MMP [35, 36]. MMP has also been preserved in various human cells lacking mitochondrial DNA [37]. Furthermore, treatment of mitochondria, derived from patients with mitochondrial dysfunction diseases, with oligomycin results in the loss of membrane potential [38]. In IF1-deficient 143C cells, co-treatment with FCCP and oligomycin decreases membrane potential [39]. It has been suggested that inhibition of the reverse mode of H^+^/ATPase in cancer cells, such as 143 B, reduces MMP [32]. In cardiac cells under ischemic or hypoxic conditions, mitochondria consume ATP through H^+^/ATPase operating in reverse mode [30, 31]. Mitochondrial ATP consumption has also been implicated in cancer and various other diseases [40, 41].

The reverse mode of H^+^/ATPase functions to maintain MMP, which is a critical aspect of cell survival since MMP is closely associated with programmed cell death [42]. Treatment with the complex I inhibitor rotenone has been reported to induce apoptosis in MCF7 [43], and loss of membrane potential can trigger apoptosis [44]. A growing body of evidence supports the notion that an unstable membrane potential amplifies apoptosis triggered by other factors [25]. Therefore, maintaining MMP plays a key role in prevention of apoptosis. Indeed, co-treatment of MCF7 cells with FCCP and oligomycin resulted in a low MMP (Fig. 4e) and induction of apoptosis (Fig. 4f).

ATP-consuming mitochondria rely on ATP transport from the cytosol to the mitochondria [45]. Moreover, Ant2 expression increases under anaerobic conditions in cancer cells such as HepG2, probably to support ATP transport from the cytosol to the mitochondria [32]. This study also showed that the *sal1*Δ*aac2*Δ double knockout strain of *S. cerevisiae* exhibited increased dead cell rate under anaerobic conditions (Fig. 4c), highlighting the critical role of ATP transport from the cytosol to the mitochondria in preventing cell death under the hypoxic condition.

Based on these results, it appears that ATP supply to mitochondria plays a key role in aerobic glycolysis, and this role is conserved in both cancer and yeast cells. This function is not always active but becomes significant when membrane potential cannot be stably maintained by the activity of electron transport chain, such as under hypoxic conditions or during unstable oxygen supply [30, 31, 33]. For example, *S. cerevisiae* is an organism specialized for niches with high sugar concentrations, such as flower nectar [46]. The high sugar and low oxygen supply environments facilitate rapid proliferation via active metabolism, which quickly depletes dissolved oxygen and creates an anaerobic micro-environment [47]. Moreover, for floating cells, the activity of the electron transport chain may become unstable owing to the fluctuating concentrations of dissolved oxygen. In such conditions, the constant activation of substrate-level phosphorylation in the cytosol provides a strategic advantage by acting as an emergency mechanism to meet sudden ATP demands from the mitochondria.

Cancer cells are survivors that resist diverse pressure from the immune system aimed at inducing cell death [48]. Cancer cells can also proliferate under hypoxic conditions with an unstable oxygen supply [40]. To endure such adverse conditions, the constant activation of substrate-level phosphorylation appears essential for maintaining the MMP at a safe level, thereby preventing the onset of apoptosis.

Although its origin remains unclear [49], aerobic glycolysis and its role in supplying ATP to mitochondria may be an evolutionarily ancient mechanism, because apoptosis is a universal mechanism in eukaryotes [50, 51]. Indeed, aerobic glycolysis and reverse-mode ATPases have been identified in various species and cell types. For example, various mammalian stem cells [52, 53] and *Drosophila* macrophages rely on aerobic glycolysis [54]. When a protist infects primary human macrophages, mitochondrial H^+^/ATPase switches to ATP hydrolase activity and prevents macrophage cell death [55]. As mentioned earlier, aerobic glycolysis plays several roles in various biological processes. A more detailed investigation of the additional roles of aerobic glycolysis will lead to a better understanding of the biological significance and clinical and industrial applications of the Warburg effect in mammalian cancer cells and the Crabtree effect in *S. cerevisiae*.

## Materials and Method

### *S. cerevisiae* strain and culture conditions

Two laboratory strains of *S. cerevisiae* S288C (BY27002) and BY18615, were purchased from the National BioResource Project (NBRP, Japan). *S. cerevisiae* strains and plasmids used in this study are listed in Table S4. *S. cerevisiae* cells were inoculated into 5 mL of yeast extract peptone dextrose (YPD) medium (1% Bacto yeast extract, 2% Bacto peptone, 2% glucose) in a test tube and cultured with shaking at 30 °C, 150 rpm for 24 h using a Personal shaker-11 (TAITEC Co. Saitama, Japan). For pre-culture, cells were transferred into 50 mL of synthetic dextrose (SD) medium (0.5% glucose, 0.67% yeast nitrogen base without amino acids) in a 200 mL baffled flask and cultured overnight with shaking at 30 °C, 120 rpm using a Bio Shaker (GBR-200, TAITEC). For the main culture, cells were inoculated into 5 mL of SD medium in a CELLSTAR 6-well plate with low evaporation lid (Greiner Bio-one, Kremsmünster, Austria) and cultured with shaking at 30 °C, 185 rpm using a Synergy HTX multi-mode plate reader (BioTek, Vermont, USA.). Cell concentration was determined by measuring the optical density at 600 nm (OD_600_) using a UV-visible spectrophotometer (UV-1280, Shimadzu, Kyoto, Japan). The initial OD_600_ values of the pre- and main cultures were 0.05.

### Cancer cell and culture conditions

The MCF7 human breast carcinoma cell line was purchased from the Riken Bioresource Center (BRC). Cells (3.6×10^5^ cells) were seeded in 5 mL of medium in 60-mm (diameter) plates and cultured at 37 °C with 5% CO_2_. For passage cultures, Dulbecco’s modified Eagle’s medium (DMEM) containing 10% fetal bovine serum (FBS) and 1% penicillin/streptomycin (Fujifilm Wako Pure Chemical Co., Osaka, Japan) was used. For metabolic analysis, cells were cultured in 5 mL DMEM without glucose, L-glutamine, phenol red, sodium pyruvate, or sodium bicarbonate (Sigma Aldrich, St. Louis, MO, USA) and supplemented with 20 mM glucose, 2 mM glutamine, 3.7 g/L sodium bicarbonate, and 10% dialyzed FBS (Lot No. 2133990, Gibco). A glucose concentration of 10 mM was employed for medium component analysis, and cell numbers were measured 10 times at any location within the dish using CKX-CCSW (OLYMPUS, Tokyo, Japan), with a culture area of 2100 mm², objective lens magnification of 10x, and adapter magnification of 0.5x.

### Measurement of extracellular metabolite concentration

For extracellular metabolite measurements, culture medium (0.1–1.0 mL) was collected, mixed with an equal volume of 20 mM pimelate solution (internal standard), and filtered through a filter cartridge (0.45-µm pore size). Extracellular metabolite concentrations were determined using high-performance liquid chromatography (HPLC; Prominence, Shimadzu) equipped with a refractive index detector (RI) and an Aminex HPX-87H column (Bio-Rad, CA, USA), as previously described [56].

### Inhibitor treatment

This study utilized three inhibitors: FCCP [Carbonyl cyanide 4-(trifluoromethoxy) phenylhydrazone] (Sigma-Aldrich), antimycin A (from *Streptomyces* sp.; Sigma-Aldrich), and oligomycin (Fujifilm Wako). For MCF7 treatments, stock solutions were prepared as 5 mM FCCP in DMSO, 100 µM antimycin A in ethanol, and 100 µM oligomycin in ethanol. For *S. cerevisiae*, the stock solutions were 5 mM FCCP in DMSO, 40 mM antimycin A in DMSO, and 40 mM oligomycin in DMSO. All stock solutions were added to the medium at a final concentration of 0.1% (v/v).

### Metabolome analysis of intracellular metabolites

*S. cerevisiae* cells (approximately OD_600_ × mL = 10) were collected from the main culture by the filtration method (PTFE membrane filter: 0.45 µm pore size, 47 mm diameter; Omnipore, Merck Millipore, Kenilworth, NJ, USA). The cells were immersed in 1.6 mL of methanol containing an internal standard (20 µM D-camphor sulfonate) and stored at -80 °C. Intracellular metabolites were extracted using the methanol/chloroform/water method with addition of 640 µL of Milli-Q water and 1.6 mL of chloroform. For MCF7 cells, the medium was quickly removed from the dish, and the cells were washed with 1 mL of phosphate-buffered saline (PBS). Immediately after washing, 800 μL of -80 °C cooled methanol was spread over the entire dish to stop metabolism. The cells were scraped on ice using a cell scraper, collected with the methanol into tubes, and stored at -80 °C. Intracellular metabolites were similarly extracted using the methanol/chloroform/water method, with 320 µL of Milli-Q water and 0.8 mL of chloroform. The resulting mixture was centrifuged (2580 ×*g*, 4 °C, 20 min), and 250 µL of the upper phase was aliquoted into six or three Eppendorf tubes, dried under vacuum using a Speed Vac (Thermo Fisher Scientific, Waltham, MA, USA). Metabolome data were obtained via ion-pair liquid chromatography-triple quadrupole mass spectrometry (LC-MS/MS) and gas chromatography-quadrupole mass spectrometry (GC-MS) were performed, as previously reported method [57]. For GC-MS analysis, dried samples were derivatized by adding 100 μL each of methoxyamine hydrochloride in pyridine (40 mg/mL) and *N*-methyl-*N*-(*tert*-butyldimethylsilyl)trifluoroacetamide (MTBSTFA) with 1% *tert*-butyldimetheylchlorosilane (TBDMCS) (Thermo Fisher). All data were processed using the LabSolution software (version 5.1, Shimadzu).

### Quantitative PCR analysis

*S. cerevisiae* cells (approximately OD_600_ × mL = 2) were collected from the main culture by centrifugation at 5000 ×*g* for 10 min. Following removal of the supernatant, the cell wall was disrupted by adding 10 μL of 1000 U/mL zymolyase (Nacalai Tesque, Kyoto, Japan) solution, 40 μL of 500 mM EDTA, and 60 μL of sterile water, followed by incubation at 37 °C for 30 min. For MCF7 cells, the medium was removed, and cells were collected following the manufacture’s protocol. RNA was extracted from the collected cells using the Nucleo SpinR RNA kit (Takara Bio, Shiga, Japan). cDNA was synthesized using the PrimeScript RT Master Mix (Perfect Real Time) kit (Takara Bio) using the TaKaRa PCR Thermal Cycler Dice (Takara Bio) (37 °C for 15 min, 85 °C for 5 sec). qPCR was conducted on a StepOnePlus Real-Time PCR System (Applied Biosystems, Waltham, MA, USA) using TB GreenR Premix Ex TaqII (Tli RNaseH Plus) (Takara Bio). Amplification conditions were Stage 1:95 °C for 5 s, Stage 2:95 °C for 5 s and 60 °C for 30 s, repeated for 40 cycles. Primers used are listed in Supplementary Table S5. Relative expression levels were calculated using the ΔΔCt method [58] using *ACT1* and *GAPDH* as internal controls for *S. cerevisiae* and MCF7 cells, respectively.

### Measurement of mitochondrial membrane potential

*S. cerevisiae* culture broth (approximately OD_600_ × mL = 1) was collected and treated with MitoTracker Red CM-H2Xros (Invitrogen, Carlsbad, CA, USA) at a final concentration of 400 nM. Following additional cultivation with shaking at 30 °C for 30 min, the medium was removed, and the cells were resuspended in 30 μL of PBS. The suspended cells were then placed on glass plates. For the analysis of MCF7 cells, 1.5 × 10^5^ cells were seeded in 1.5 mL of passage medium in a 35 mm glass-bottom dish (Matsunami Glass Ind., Osaka, Japan). After overnight cultivation, the medium was replaced with metabolic analysis medium containing 0.1% (v/v) inhibitors. Following 3 h of culture, 750 μL of supernatant was removed, and 750 μL of MitoTracker Red CM-H2Xros (400 nM, diluted in the metabolic analysis medium). After 30 min of incubation, the medium was removed and replaced with 1 mL of PBS. Fluorescence microscopy observation was performed using an IX71 Fluorescence Microscope (Olympus) equipped with a U-MWIG3 filter (Ex530-550/DM570/BA575-625, Olympus) and an ORCA-spark digital CMOS camera C11578-36U (Hamamatsu Photonics, Aichi, Japan). All images were processed and analyzed using CellSens (Olympus) and Fiji [59].

### Measurement of mitochondrial membrane potential under hypoxic conditions

For *S. cerevisiae*, cells in pre-culture were suspended in 1 mL SD medium with OD_600_ = 0.05 in 1.5 mL microcentrifuge tubes (Eppendorf) and incubated at 30 °C for 24 h in an AnaeroPack rectangular jar standard type (Sugiyama-Gen Co. Tokyo, Japan) containing an AnaeroPack-Kenki oxygen scavenger (Sugiyama-Gen). Following treatment with MitoTracker Red CM-H2Xros at a final concentration of 400 nM, the cell suspension was transferred to a 35 mm glass-bottom dish (AGC Techno glass, Co, Shizuoka, Japan) and cultured for 30 min in a low-oxygen incubator CH-070A (BLAST, Kanagawa, Japan). Fluorescence observations were made with the dish remaining in the culture chamber.

For MCF7 cells (1.5 × 10^5^ cells) were seeded in 1.5 mL of passage medium in a 35 mm glass-bottom dish and cultured overnight to allow attachment of the cells to the bottom of the dish. The medium was replaced with the metabolic analysis medium containing 0.1% (v/v) inhibitors and cultured at 5% CO_2_, 1% O_2_, 37 °C for 24 h using a low-oxygen incubator CH-070A. An AnaeroPack-Kenki was used as an oxygen scavenger. After treatment with MitoTracker Red CM-H2Xros,as described above, fluorescence observations were conducted while maintaining the hypoxic state by keeping the dish in the low-oxygen incubator.

### Construction of *S. cerevisiae* strains expressing cytoplasmic and mitochondria-localized QUEEN-2m

Based on the amino acid sequence of the intracellular ATP sensor QUEEN-2m [26], a codon-optimized ORF was synthesized and cloned into the pGEM-T Easy vector (Promega, Madison, WI, USA). The pGEM-T Easy-QUEEN-2m plasmid and a multicopy yeast expression plasmid pGK426 [60], were digested with SalI and EcoRI and the resulting fragments were ligated to construct pGK426-QUEEN-2m.

To create the mitochondria-localized QUEEN-2m expression plasmid (pAS2, Figure S6), inverse PCR was performed using the pGK426-QUEEN-2m as the template. Primers (Queen-2m_Cox4N_invF and Queen-2m_Cox4N_invR; Table S6) were designed to amplify outward from the N-terminus of the QUEEN-2m gene, including the sequence encoding the mitochondrial targeting signal derived from *S. cerevisiae* Cox4. The obtained linear DNA fragment was phosphorylated using T4 Polynucleotide Kinase (New England Labs, Ipswich, MA, USA) and ligated using T4 DNA Ligase and T4 Polynucleotide Kinase (New England Labs) with T4 DNA Ligase (New England Labs). The circularized plasmids were transformed into *Escherichia. coli* DH5α, and plated on LB agar containing ampicillin (100 μg/mL). Plasmid DNA was extracted from colonies grown on the selection plates, and the presence of the correctly introduced Cox4 pre-sequence was confirming by sequencing. The construct was designed as to create a circularized plasmid pAS1.

Furthermore, to construct pAS2, which harbors two ORFs of QUEEN-2m, inverse PCR was performed on pAS1 using primers (AS1 fusion-F1 and AS1 fusion-R1; Table S6) starting 670 bp upstream of the PGK1 terminator. An additional ORF was obtained by PCR using pAS1 as a template and primers AS1 QUEEN-F1 and AS1 QUEEN-R1 (Table S6). The obtained vector and fragment were fused using the Gibson Assembly method with the Gibson Assembly Master Mix (New England Biolabs, Ipswich, MA, USA) and transformed into *E. coli* DH5α. Plasmid DNA was extracted from colonies grown on LB agar containing ampicillin (100 µg/mL), and sequencing confirmed the presence of two copies of the QUEEN-2m ORFs in the construct. The *S. cerevisiae* strain BY18615 (MATα, ura3) (purchased from the National BioResource Project, NBRP) was transformed with the pGK426-QUEEN-2m and pAS2 plasmids using the standard. The lithium acetate method was used for transformation method.

### Measurement of ATP concentration in cytoplasm and mitochondria using QUEEN-2m expressed *S. cerevisiae* strains

The analysis was performed following previously described protocol [61]. *S. cerevisiae* cells were anchored onto glass plates coated with concanavalin A. A 35 mm glass-bottom dish was prepared by soaking with 60 μL of concanavalin A solution (2 mg/mL, Sigma) for 5 min. The concanavalin A solution was then removed, and the dish was washed twice with 100 μL of sterile water. From a log-phase pre-culture of *S. cerevisiae* strain expressing QUEEN-2m, 100 μL of culture broth was collected and placed on the prepared dish for 5 min. After carefully removing of 80 μL of the medium to avoid detaching the cells, 80 μL of fresh SD medium was added, gently suspended five times, and removed. This process was repeated twice, followed by addition of 80 μL of fresh SD medium.

Fluorescence observations were made using an inverted fluorescence microscope (ECLIPSE TE2000-E, Nikon, Tokyo, Japan) equipped with a 60× oil immersion objective lens (Plan Apo 60×1.40 Oil Ph3 DM), a GFP filter set (Ex457-481/DM495/BA502-538, Nikon), a QUEEN filter set (Ex393-425/DM506/BA516-556, Semrock), and an EM-CCD camera IXON (Nikon). During fluorescence imaging, the sample was maintained at 30 °C using a microscope temperature control system (TOKAI HIT, Shizuoka, Japan). To minimize sample fading, the exposure time was set to 50 ms for both GFP and QUEEN fluorescence. Images were captured every 10 min, and the response after inhibitor treatment was observed continuously for 120 min.

The fluorescence image data were analyzed using Fiji [59]. All images were converted to 32-bit format after background subtraction using the rolling-ball algorithm. A threshold level for detecting cell regions was set using the IsoData algorithm, with pixel values outside the detected cell regions assigned as NaN values. The QUEEN/GFP ratio was determined using processed fluorescence images of GFP and QUEEN. For each captured image, cell regions were extracted, and the mean intensity ratio was calculated for each cell type. For time-course data, stacks were created for both GFP and QUEEN fluorescence images, and image processing was performed for each stack. The LPixel plugin was used to correct any image shifts during continuous shooting.

## Supporting information

Supplementary Fgireus and Tables

## Author Contributions

Conceptualization, A. S., M. T., and F. M.; validation and investigation, A. S., T. T., K. N., T. S., and N. O.; writing—original draft preparation, A. S., N. O., and F. M.; writing—review and editing, N. O., M. T., and F. M.

## Data Availability Statement

The data presented in this study are available upon request from the corresponding author.

## Acknowledgments

We thank Prof. Junko Iida, Yuki Ito, Akihiro Kunisawa, from Shimadzu Corporation for their helpful comments and technical assistance. The plasmid vector (pGK426) was provided by the National Unk Resource Project (NBRP), Japan.

## Supplementary figures and tables

**Figure S1** Fluorescent imaging of mitochondrial membrane potential using MitoTracker reagent.

**Figure S2** Fluorescent imaging of mitochondrial membrane potential of inhibitor treated human breast cancer cell, MCF7 and *S. cerevisiae* S288C.

**Figure S3** Localized expression of QUEEN-2m in cytosol and mitochondria.

**Figure S4** Fluorescent imaging of mitochondrial membrane potential of human breast cancer cell, MCF7 and *S. cerevisiae* S288C treated with FCCP and oligomycin.

**Figure S5** Fluorescent imaging of mitochondrial membrane potential of human breast cancer cell, MCF7 and *S. cerevisiae* S288C treated with oligomycin under hypoxic conditions.

**Figure S6** Vector map of pAS2

**Table S1** Specific rates for cell proliferation, glucose uptake, ethanol and glycerol production of inhibitor treated *S. cerevisiae* (S288C) cells during log-phase.

**Table S2** Specific rates for cell proliferation, glucose uptake, and lactate production of inhibitor treated human breast cancer (MCF7) cells during log-phase.

**Table S3** Metabolic profile data obtained from inhibitor treated S288C and MCF7 cells

**Table S4** *Saccharomyces cerevisiae* strains and plasmids used in this study

**Table S5** Primers for qPCR

**Table S6** Primers to construct pAS1 and pAS2

## References

1. Diaz-Ruiz R, Rigoulet M, Devin A: The Warburg and Crabtree effects: On the origin of cancer cell energy metabolism and of yeast glucose repression. Biochim Biophys Acta 2011, 1807(6):568–576.

2. Vaupel P, Multhoff G: Revisiting the Warburg effect: historical dogma versus current understanding. J Physiol 2021, 599(6):1745–1757.

3. Pavlova NN, Thompson CB: The emerging hallmarks of cancer metabolism. Cell Metab 2016, 23(1):27–47.

4. Zu XL, Guppy M: Cancer metabolism: facts, fantasy, and fiction. Biochem Biophys Res Commun 2004, 313(3):459–465.

5. Moreno-Sanchez R, Rodriguez-Enriquez S, Marin-Hernandez A, Saavedra E: Energy metabolism in tumor cells. FEBS J 2007, 274(6):1393–1418.

6. Tran Q, Lee H, Park J, Kim SH, Park J: Targeting cancer metabolism - eevisiting the Warburg effects. Toxicol Res 2016, 32(3):177–193.

7. Pfeiffer T, Morley A: An evolutionary perspective on the Crabtree effect. Front Mol Biosci 2014, 1:17.

8. Hammad N, Rosas-Lemus M, Uribe-Carvajal S, Rigoulet M, Devin A: The Crabtree and Warburg effects: Do metabolite-induced regulations participate in their induction? Biochim Biophys Acta 2016, 1857(8):1139–1146.

9. Lunt SY, Vander Heiden MG: Aerobic glycolysis: meeting the metabolic requirements of cell proliferation. Annu Rev Cell Dev Biol 2011, 27:441–464.

10. Liberti MV, Locasale JW: The Warburg effect: How does it benefit cancer cells? Trends Biochem Sci 2016, 41(3):211–218.

11. Niebel B, Leupold S, Heinemann M: An upper limit on Gibbs energy dissipation governs cellular metabolism. Nat Metab 2019, 1(1):125–132.

12. Martins Pinto M, Paumard P, Bouchez C, Ransac S, Duvezin-Caubet S, Mazat JP, Rigoulet M, Devin A: The Warburg effect and mitochondrial oxidative phosphorylation: Friends or foes? Biochim Biophys Acta Bioenerg 2023, 1864(1):148931.

13. Cerulus B, Jariani A, Perez-Samper G, Vermeersch L, Pietsch JMJ, Crane MM, New AM, Gallone B, Roncoroni M, Dzialo MC et al: Transition between fermentation and respiration determines history-dependent behavior in fluctuating carbon sources. Elife 2018, 7.

14. Chaube B, Malvi P, Singh SV, Mohammad N, Meena AS, Bhat MK: Targeting metabolic flexibility by simultaneously inhibiting respiratory complex I and lactate generation retards melanoma progression. Oncotarget 2015, 6(35):37281–37299.

15. Vemuri GN, Eiteman MA, McEwen JE, Olsson L, Nielsen J: Increasing NADH oxidation reduces overflow metabolism in *Saccharomyces cerevisiae*. Proc Natl Acad Sci U S A 2007, 104(7):2402–2407.

16. van Maris AJA, Bakker BM, Brandt M, Boorsma A, de Mattos MJT, Grivell LA, Pronk JT, Blom J: Modulating the distribution of fluxes among respiration and fermentation by overexpression of HAP4 in *Saccharomyces cerevisiae*. FEMS Yeast Res 2001, 1(2):139–149.

17. Araki C, Okahashi N, Maeda K, Shimizu H, Matsuda F: Mass spectrometry-based method to study inhibitor-induced metabolic redirection in the central metabolism of cancer cells. Mass Spectrometry 2018, 7(1):A0067.

18. Kitajima S, Yoshida A, Kohno S, Li F, Suzuki S, Nagatani N, Nishimoto Y, Sasaki N, Muranaka H, Wan Y et al: The RB-IL-6 axis controls self-renewal and endocrine therapy resistance by fine-tuning mitochondrial activity. Oncogene 2017, 36(36):5145–5157.

19. Wu L, van Dam J, Schipper D, Kresnowati MT, Proell AM, Ras C, van Winden WA, van Gulik WM, Heijnen JJ: Short-term metabolome dynamics and carbon, electron, and ATP balances in chemostat-grown *Saccharomyces cerevisiae* CEN.PK 113-7D following a glucose pulse. Appl Environ Microbiol 2006, 72(5):3566–3577.

20. Theobald U, Mailinger W, Baltes M, Rizzi M, Reuss M: In vivo analysis of metabolic dynamics in Saccharomyces cerevisiae : I. Experimental observations. Biotechnol Bioeng 1997, 55(2):305–316.

21. Kresnowati MT, van Winden WA, Almering MJ, ten Pierick A, Ras C, Knijnenburg TA, Daran-Lapujade P, Pronk JT, Heijnen JJ, Daran JM: When transcriptome meets metabolome: fast cellular responses of yeast to sudden relief of glucose limitation. Mol Syst Biol 2006, 2:49.

22. Ganapathy V, Thangaraju M, Prasad PD: Nutrient transporters in cancer: relevance to Warburg hypothesis and beyond. Pharmacol Ther 2009, 121(1):29–40.

23. Ozcan S, Johnston M: Three different regulatory mechanisms enable yeast hexose transporter (HXT) genes to be induced by different levels of glucose. Mol Cell Biol 1995, 15(3):1564–1572.

24. Galardo MN, Riera MF, Pellizzari EH, Cigorraga SB, Meroni SB: The AMP-activated protein kinase activator, 5-aminoimidazole-4-carboxamide-1-b-D-ribonucleoside, regulates lactate production in rat Sertoli cells. J Mol Endocrinol 2007, 39(4):279–288.

25. Gottlieb E, Armour SM, Harris MH, Thompson CB: Mitochondrial membrane potential regulates matrix configuration and cytochrome c release during apoptosis. Cell Death Differ 2003, 10(6):709–717.

26. Yaginuma H, Kawai S, Tabata KV, Tomiyama K, Kakizuka A, Komatsuzaki T, Noji H, Imamura H: Diversity in ATP concentrations in a single bacterial cell population revealed by quantitative single-cell imaging. Sci Rep 2014, 4:6522.

27. Kubota T, Shindo Y, Tokuno K, Komatsu H, Ogawa H, Kudo S, Kitamura Y, Suzuki K, Oka K: Mitochondria are intracellular magnesium stores: investigation by simultaneous fluorescent imagings in PC12 cells. Biochim Biophys Acta 2005, 1744(1):19–28.

28. Rimessi A, Giorgi C, Pinton P, Rizzuto R: The versatility of mitochondrial calcium signals: from stimulation of cell metabolism to induction of cell death. Biochim Biophys Acta 2008, 1777(7-8):808–816.

29. Acin-Perez R, Beninca C, Fernandez Del Rio L, Shu C, Baghdasarian S, Zanette V, Gerle C, Jiko C, Khairallah R, Khan S et al: Inhibition of ATP synthase reverse activity restores energy homeostasis in mitochondrial pathologies. EMBO J 2023, 42(10):e111699.

30. Rouslin W, Broge CW: Mechanisms of ATP conservation during ischemia in slow and fast heart rate hearts. Am J Physiol 1993, 264(1 Pt 1):C209–216.

31. Solaini G, Harris DA: Biochemical dysfunction in heart mitochondria exposed to ischaemia and reperfusion. Biochem J 2005, 390(Pt 2):377–394.

32. Chevrollier A, Loiseau D, Gautier F, Malthiery Y, Stepien G: ANT2 expression under hypoxic conditions produces opposite cell-cycle behavior in 143B and HepG2 cancer cells. Mol Carcinog 2005, 42(1):1–8.

33. Traba J, Froschauer EM, Wiesenberger G, Satrustegui J, Del Arco A: Yeast mitochondria import ATP through the calcium-dependent ATP-Mg/Pi carrier Sal1p, and are ATP consumers during aerobic growth in glucose. Mol Microbiol 2008, 69(3):570–585.

34. Peeters K, Van Leemputte F, Fischer B, Bonini BM, Quezada H, Tsytlonok M, Haesen D, Vanthienen W, Bernardes N, Gonzalez-Blas CB et al: Fructose-1,6-bisphosphate couples glycolytic flux to activation of Ras. Nat Commun 2017, 8(1):922.

35. Chinopoulos C, Tretter L, Adam-Vizi V: Depolarization of in situ mitochondria due to hydrogen peroxide-induced oxidative stress in nerve terminals:: Inhibition of α-ketoglutarate dehydrogenase. J Neurochem 1999, 73(1):220–228.

36. Scott ID, Nicholls DG: Energy transduction in intact synaptosomes. Influence of plasma-membrane depolarization on the respiration and membrane potential of internal mitochondria determined in situ. Biochem J 1980, 186(1):21–33.

37. Appleby RD, Porteous WK, Hughes G, James AM, Shannon D, Wei YH, Murphy MP: Quantitation and origin of the mitochondrial membrane potential in human cells lacking mitochondrial DNA. Eur J Biochem 1999, 262(1):108–116.

38. McKenzie M, Liolitsa D, Akinshina N, Campanella M, Sisodiya S, Hargreaves I, Nirmalananthan N, Sweeney MG, Abou-Sleiman PM, Wood NW et al: Mitochondrial ND5 gene variation associated with encephalomyopathy and mitochondrial ATP consumption. J Biol Chem 2007, 282(51):36845–36852.

39. Sgarbi G, Barbato S, Costanzini A, Solaini G, Baracca A: The role of the ATPase inhibitor factor 1 (IF(1)) in cancer cells adaptation to hypoxia and anoxia. Biochim Biophys Acta Bioenerg 2018, 1859(2):99–109.

40. Solaini G, Baracca A, Lenaz G, Sgarbi G: Hypoxia and mitochondrial oxidative metabolism. Biochim Biophys Acta 2010, 1797(6-7):1171–1177.

41. Galber C, Acosta MJ, Minervini G, Giorgio V: The role of mitochondrial ATP synthase in cancer. Biol Chem 2020, 401(11):1199–1214.

42. Kroemer G, Galluzzi L, Brenner C: Mitochondrial membrane permeabilization in cell death. Physiol Rev 2007, 87(1):99–163.

43. Deng YT, Huang HC, Lin JK: Rotenone induces apoptosis in MCF-7 human breast cancer cell-mediated ROS through JNK and p38 signaling. Mol Carcinog 2010, 49(2):141–151.

44. McClintock DS, Santore MT, Lee VY, Brunelle J, Budinger GR, Zong WX, Thompson CB, Hay N, Chandel NS: Bcl-2 family members and functional electron transport chain regulate oxygen deprivation-induced cell death. Mol Cell Biol 2002, 22(1):94–104.

45. Zorova LD, Popkov VA, Plotnikov EY, Silachev DN, Pevzner IB, Jankauskas SS, Babenko VA, Zorov SD, Balakireva AV, Juhaszova M et al: Mitochondrial membrane potential. Anal Biochem 2018, 552:50–59.

46. Marsit S, Leducq JB, Durand E, Marchant A, Filteau M, Landry CR: Evolutionary biology through the lens of budding yeast comparative genomics. Nat Rev Genet 2017, 18(10):581–598.

47. Enfors SO, Jahic M, Rozkov A, Xu B, Hecker M, Jurgen B, Kruger E, Schweder T, Hamer G, O’Beirne D et al: Physiological responses to mixing in large scale bioreactors. J Biotechnol 2001, 85(2):175–185.

48. Waldman AD, Fritz JM, Lenardo MJ: A guide to cancer immunotherapy: from T cell basic science to clinical practice. Nat Rev Immunol 2020, 20(11):651–668.

49. Rio Bartulos C, Rogers MB, Williams TA, Gentekaki E, Brinkmann H, Cerff R, Liaud MF, Hehl AB, Yarlett NR, Gruber A et al: Mitochondrial glycolysis in a major lineage of eukaryotes. Genome Biol Evol 2018, 10(9):2310–2325.

50. Koonin EV, Aravind L: Origin and evolution of eukaryotic apoptosis: the bacterial connection. Cell Death Differ 2002, 9(4):394–404.

51. Ameisen JC: On the origin, evolution, and nature of programmed cell death: a timeline of four billion years. Cell Death Differ 2002, 9(4):367–393.

52. Simon MC, Keith B: The role of oxygen availability in embryonic development and stem cell function. Nat Rev Mol Cell Biol 2008, 9(4):285–296.

53. Ito K, Suda T: Metabolic requirements for the maintenance of self-renewing stem cells. Nat Rev Mol Cell Biol 2014, 15(4):243–256.

54. Krejcova G, Danielova A, Nedbalova P, Kazek M, Strych L, Chawla G, Tennessen JM, Lieskovska J, Jindra M, Dolezal T et al: Drosophila macrophages switch to aerobic glycolysis to mount effective antibacterial defense. Elife 2019, 8.

55. Escoll P, Platon L, Drame M, Sahr T, Schmidt S, Rusniok C, Buchrieser C: Reverting the mode of action of the mitochondrial F_O_F_1_-ATPase by *Legionella pneumophila* preserves its replication niche. Elife 2021, 10.

56. Ishii J, Morita K, Ida K, Kato H, Kinoshita S, Hataya S, Shimizu H, Kondo A, Matsuda F: A pyruvate carbon flux tugging strategy for increasing 2,3-butanediol production and reducing ethanol subgeneration in the yeast *Saccharomyces cerevisiae*. Biotechnol Biofuels 2018, 11:180.

57. Nishiguchi H, Liao J, Shimizu H, Matsuda F: Novel allosteric inhibition of phosphoribulokinase identified by ensemble kinetic modeling of Synechocystis sp. PCC 6803 metabolism. Metab Eng Commun 2020, 11:e00153.

58. Livak KJ, Schmittgen TD: Analysis of relative gene expression data using real-time quantitative PCR and the 2^-ΔΔCT^ Method. Methods 2001, 25(4):402–408.

59. Schindelin J, Arganda-Carreras I, Frise E, Kaynig V, Longair M, Pietzsch T, Preibisch S, Rueden C, Saalfeld S, Schmid B et al: Fiji: an open-source platform for biological-image analysis. Nat Methods 2012, 9(7):676–682.

60. Ishii J, Izawa K, Matsumura S, Wakamura K, Tanino T, Tanaka T, Ogino C, Fukuda H, Kondo A: A simple and immediate method for simultaneously evaluating expression level and plasmid maintenance in yeast. J Biochem 2009, 145(6):701–708.

61. Takaine M, Ueno M, Kitamura K, Imamura H, Yoshida S: Reliable imaging of ATP in living budding and fission yeast. J Cell Sci 2019, 132(8).

