## Supplementary Fgireus and Tables for "Conserved metabolic adaptation mechanisms in cancer cells and yeast against mitochondrial dysfunction indicate an additional role of aerobic glycolysis for cell survival"

**Figure S1** Fluorescent imaging of mitochondrial membrane potential using MitoTracker reagent.

**Figure S2** Fluorescent imaging of mitochondrial membrane potential of inhibitor treated human breast cancer cell, MCF7 and *S. cerevisiae* S288C.

**Figure S3** Localized expression of QUEEN-2m in cytosol and mitochondria.

**Figure S4** Fluorescent imaging of mitochondrial membrane potential of human breast cancer cell, MCF7 and *S. cerevisiae* S288C treated with FCCP and oligomycin.

**Figure S5** Fluorescent imaging of mitochondrial membrane potential of human breast cancer cell, MCF7 and *S. cerevisiae* S288C treated with oligomycin under hypoxic conditions.

**Figure S6** Vector map of pAS2

**Table S1** Specific rates for cell proliferation, glucose uptake, ethanol and glycerol production of inhibitor treated *S. cerevisiae* (S288C) cells during log-phase.

**Table S2** Specific rates for cell proliferation, glucose uptake, and lactate production of inhibitor treated human breast cancer (MCF7) cells during log-phase.

**Table S3** Metabolic profile data obtained from inhibitor treated S288C and MCF7 cells

**Table S4** *Saccharomyces cerevisiae* strains and plasmids used in this study

**Table S5** Primers for qPCR

**Table S6** Primers to construct pAS1 and pAS2

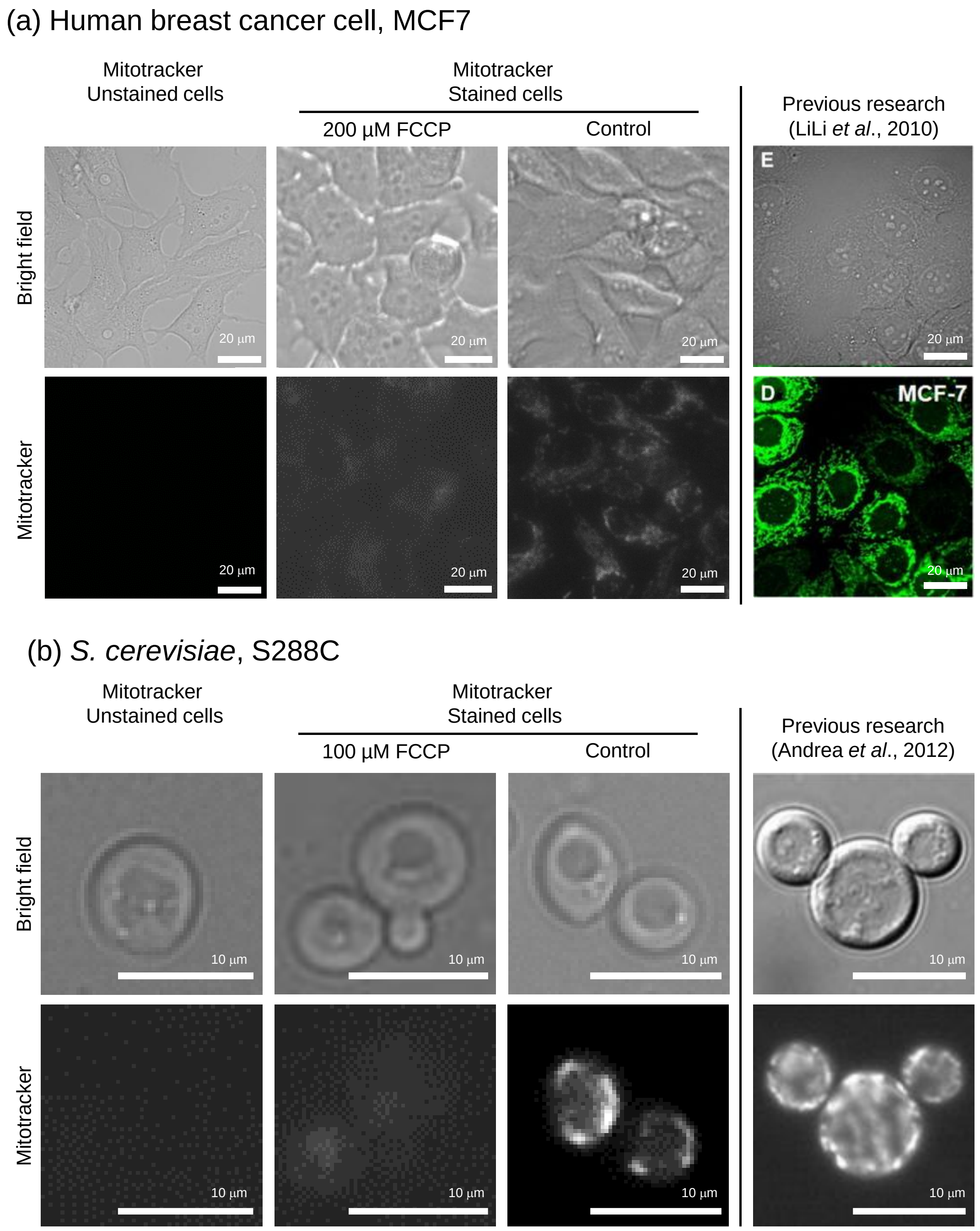


**Figure S1 Fluorescent imaging of mitochondrial membrane potential using MitoTracker reagent.** MitoTracker reagent was treated to (a) MCF-7 and (b) S288C at 3 and 2 hours after inhibitor treatment, respectively, and served for the fluorescent microscopy.

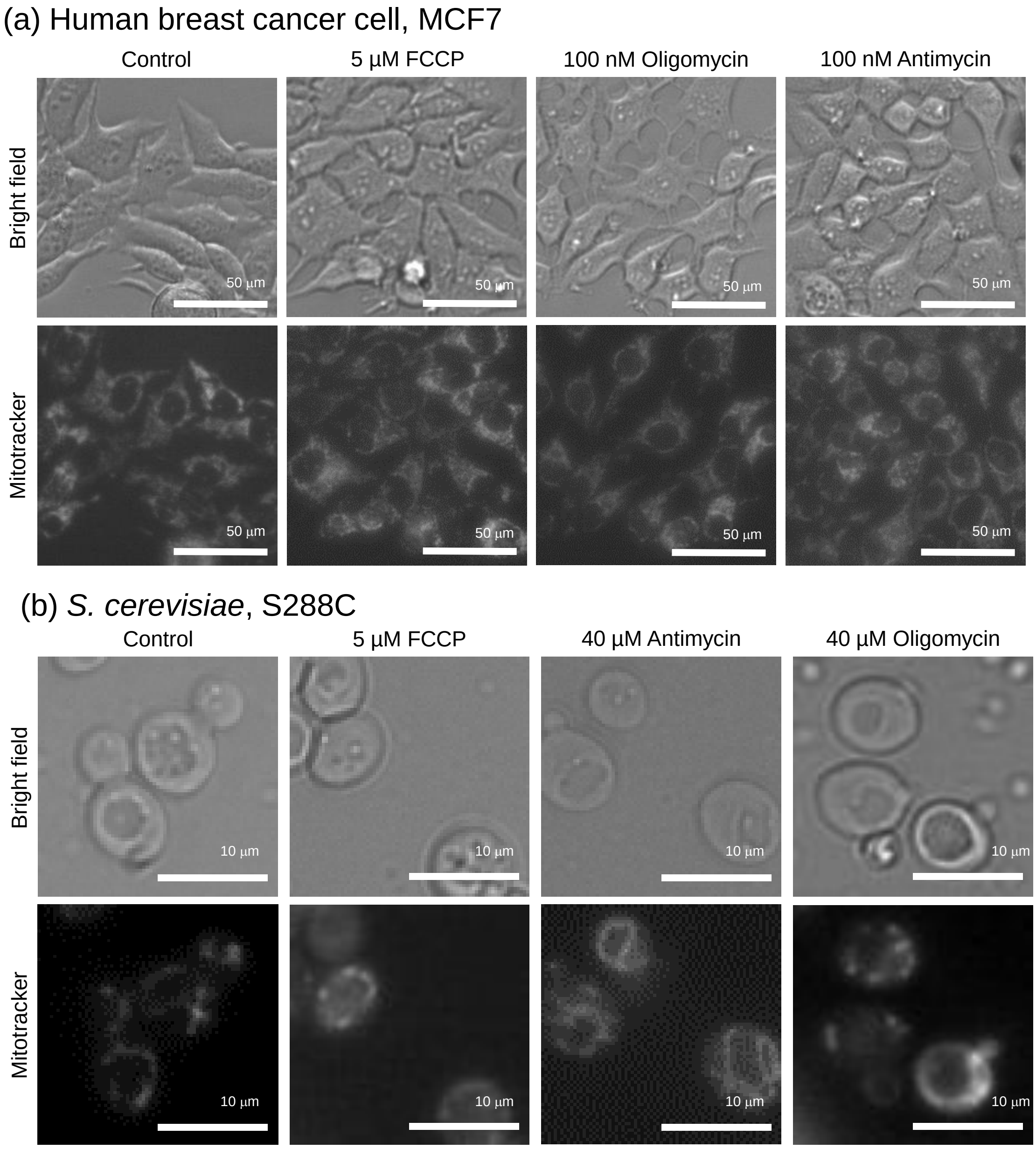


**Figure S2 Fluorescent imaging of mitochondrial membrane potential of inhibitor treated human breast cancer cell, MCF7 and *S. cerevisiae* S288C.** MitoTracker reagent was treated to (a) MCF-7 and (b) S288C cells at 3 and 2 hours after FCCP, antimycin, and oligomycin treatment, respectively, and served for the fluorescent microscopy.

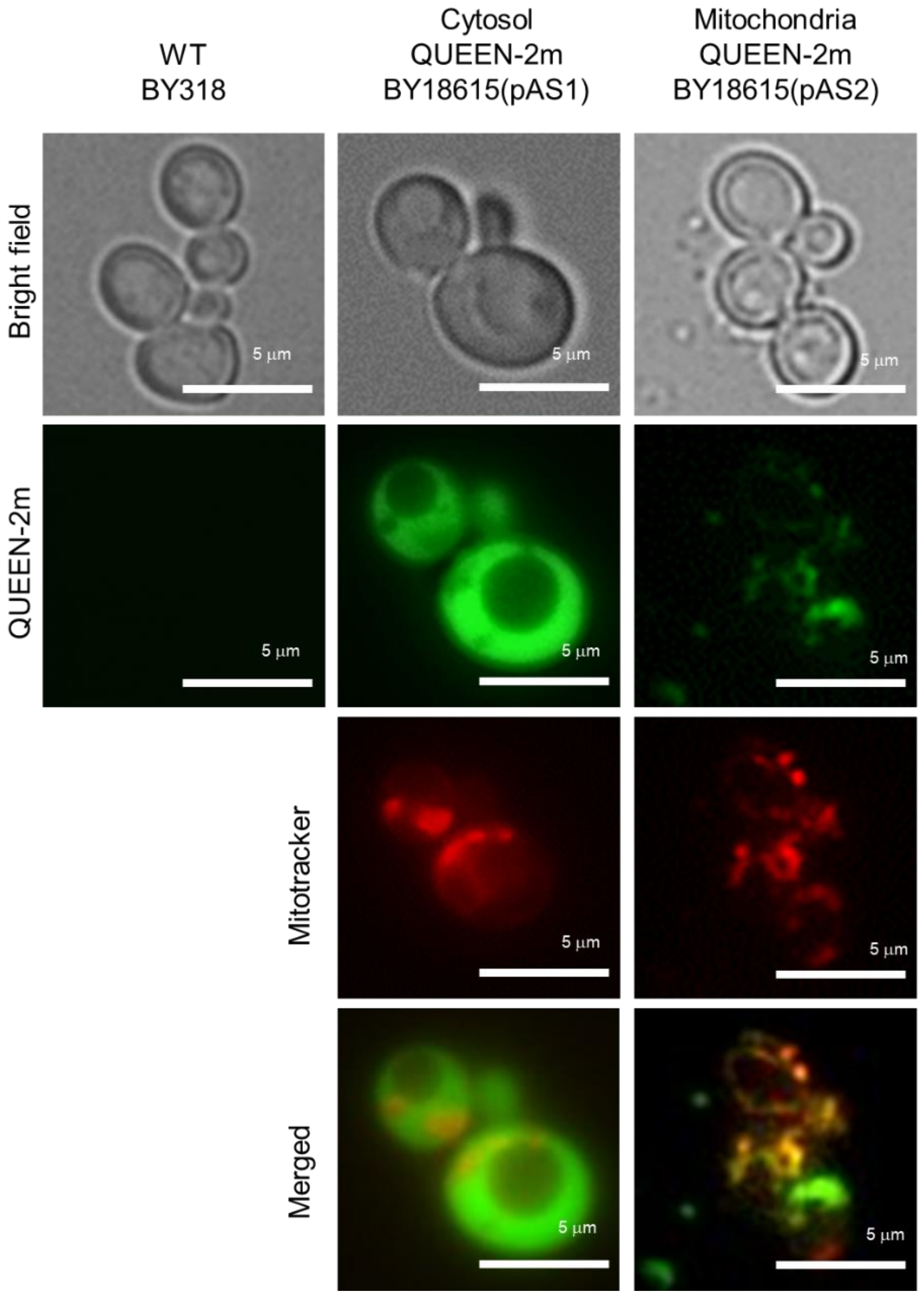


**Figure S3 Localized expression of QUEEN-2m in cytosol and mitochondria of *S. cerevisiae* S288C.** Images obtained using blight light, a green fluorescence filter, and a red fluorescence filter are shown.

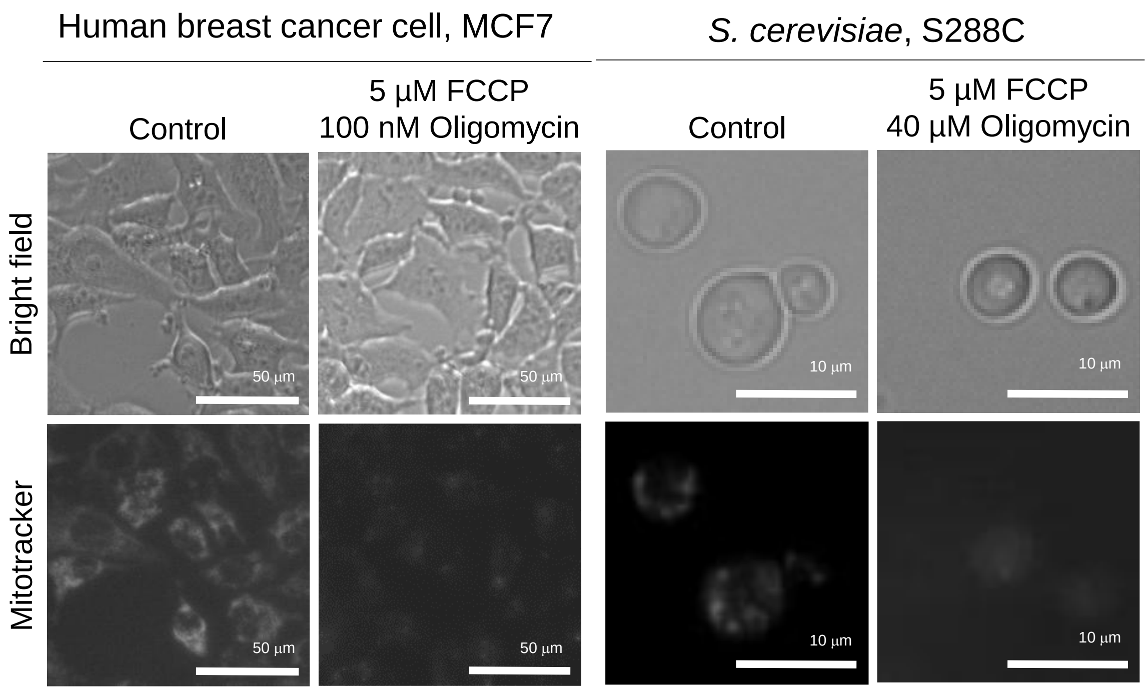


**Figure S4 Fluorescent imaging of mitochondrial membrane potential of human breast cancer cell, MCF7 and *S. cerevisiae* S288C treated with FCCP and oligomycin.** MitoTracker reagent was treated to (a) MCF-7 and (b) S288C at 3 and 2 hours after the co-treatment of FCCP and oligomycin, respectively, and served for the fluorescent microscopy.

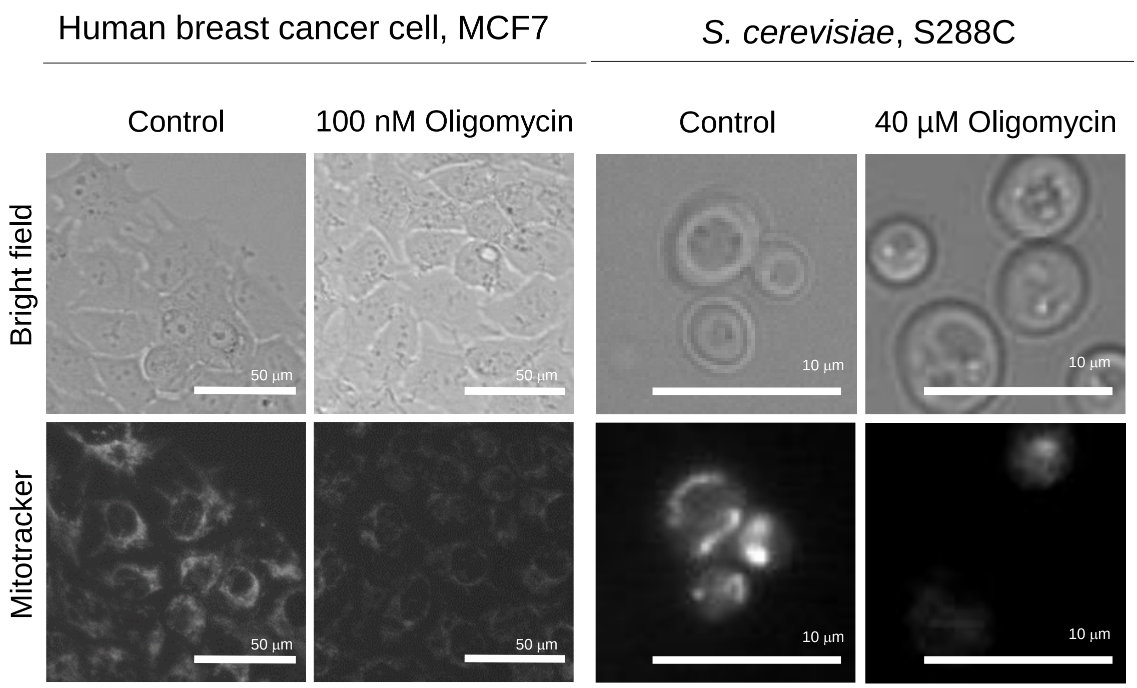


**Figure S5 Fluorescent imaging of mitochondrial membrane potential of human breast cancer cell, MCF7 and *S. cerevisiae* S288C treated with oligomycin under hypoxic conditions.** MitoTracker reagent was treated to (a) MCF-7 and (b) S288C cultured under low oxygen conditions with oligomycin and served for fluorescent microscopy.

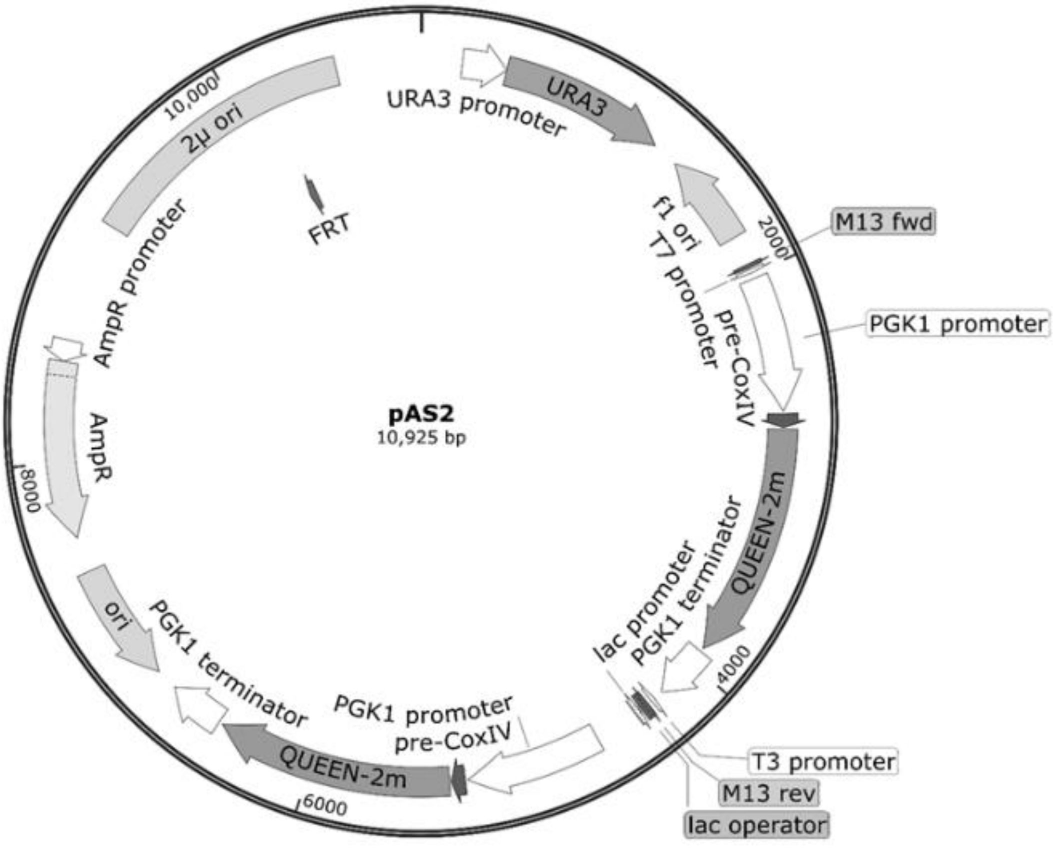


**Figure S6** Vector map of pAS2

**Table S1** Specific rates for cell proliferation, glucose uptake, ethanol and glycerol production of inhibitor treated *S. cerevisiae* (S288C) cells during log-phase.

| Inhibitors | Specific rates | | | |
|---|---|---|---|---|
| | Cell proliferation ($h^{-1}$) | Glucose uptake (mmol (gDCW h)$^{-1}$) | Ethanol production (mmol (gDCW h)$^{-1}$) | Glycerol production (mmol (gDCW h)$^{-1}$) |
| Control | 0.363±0.011 | 14.72±0.52 | 20.71±0.76 | 0.17±0.01 |
| 5 µM FCCP | 0.328±0.005* | 21.02±0.39* | 31.08±0.42* | 0.18±0.00 |
| 40 µM antimycin | 0.312±0.012* | 18.07±1.40* | 29.13±2.73* | 0.34±0.03* |
| 40 µM oligomycin | 0.315±0.010* | 17.72±1.21* | 28.90±1.86* | 0.26±0.02* |

The specific rates were determined from the growth curve. All results were obtained from triplicate cultures and described as Mean ±SD. Asterisks indicate p-value < 0.05 by two-sided t-test.

DCW: dry cell weight

**Table S2** Specific rates for cell proliferation, glucose uptake, and lactate production of inhibitor treated human breast cancer (MCF7) cells during log-phase.

| | Specific rates | | |
|---|---|---|---|
| Inhibitors | Cell proliferation ($h^{-1}$) | Glucose uptake (nmol $(10^6$cells h$)^{-1}$) | Lactate production (nmol $(10^6$cells h$)^{-1}$) |
| Control | 0.0349±0.0017 | 1237±28 | 2053±21 |
| 5 μM FCCP | 0.0194±0.0015* | 1822±250* | 3114±222* |
| 100 nM antimycin | 0.0201±0.0032* | 1696±91* | 3188±170* |
| 100 nM oligomycin | 0.0191±0.0059* | 1756±124* | 3316±184* |

The specific rates were determined from the growth curve. All results were obtained from triplicate cultures and described as Mean ±SD. Asterisks indicate p-value < 0.05 by two-sided t-test.

**Table S3** Metabolic profile data obtained from inhibitor treated S288C and MCF7 cells

| | *S. cerevisiae,* S288C | | | Human breast cancer cell, MCF7 | | |
|---|---|---|---|---|---|---|
| | Log2(Treated/Control) | | | | | |
| | FCCP | Antimycin A | Oligomycin | FCCP | Antimycin A | Oligomycin |
| G6P | 0.333 | 0.270 | 0.156 | 0.381 | -0.070 | -0.176 |
| F6P | 0.183 | 0.268 | 0.198 | 0.700 | 0.825 | -0.051 |
| FBP | 0.660 | 0.278 | -0.037 | 0.498 | 0.704 | 0.772 |
| DHAP | 0.437 | 0.511 | 0.450 | 0.696 | 0.635 | 0.394 |
| GAP | 0.292 | 0.558 | 0.832 | 0.482 | 0.514 | 0.273 |
| 3PG+2PG | -0.363 | -0.175 | -0.234 | -0.343 | -0.117 | -0.259 |
| PEP | -0.061 | -0.402 | -0.382 | -0.133 | -0.558 | -0.077 |
| Pyr | -0.125 | -0.420 | -0.739 | -0.432 | -0.972 | -0.585 |
| Lactate | nd | nd | nd | 0.461 | 0.703 | 0.265 |
| Citrate | 0.452 | -0.257 | -0.233 | 0.230 | -1.271 | -1.046 |
| 2KG | 0.077 | -0.310 | -0.108 | 0.118 | -0.826 | -0.400 |
| Succinate | 0.545 | 0.863 | 0.075 | 1.396 | 1.247 | -0.900 |
| Fumarate | -0.426 | 0.401 | 0.647 | -0.500 | 0.832 | 0.790 |
| Malate | 0.014 | 0.345 | 0.773 | -0.131 | 0.622 | 0.900 |
| 6PG | -0.845 | -0.134 | 0.032 | -1.016 | -0.741 | -0.346 |
| R5P | 0.452 | 0.508 | -0.337 | 0.162 | 0.326 | -0.006 |
| Ru5P | -0.197 | 0.120 | -0.192 | -0.217 | 0.316 | -0.191 |
| S7P | -0.471 | -0.362 | -0.216 | -0.022 | -0.262 | -0.501 |
| ATP | -0.182 | -0.148 | -0.152 | -0.060 | -0.235 | -0.067 |
| ADP | 0.069 | 0.158 | 0.034 | 0.156 | 0.359 | -0.012 |
| AMP | 0.337 | 0.783 | 0.399 | 0.482 | 0.390 | 0.254 |
| NAD | 0.195 | 0.012 | 0.398 | 0.880 | 0.207 | 0.650 |
| NADH | -0.718 | -0.811 | 0.785 | 0.222 | -2.453 | 0.327 |
| ATP/ADP | -0.294 | -0.293 | -0.176 | -0.215 | -0.583 | -0.059 |
| ATP/AMP | -0.525 | -0.870 | -0.522 | -0.545 | -0.626 | -0.319 |
| NADH/NAD | -0.901 | -0.856 | 0.385 | -0.660 | -2.664 | -0.327 |

| | *S. cerevisiae,* S288C | | | Human breast cancer cell, MCF7 | | |
|---|---|---|---|---|---|---|
| | Two-sided t test (p-value) | | | | | |
| | FCCP | Antimycin A | Oligomycin | FCCP | Antimycin A | Oligomycin |
| G6P | 0.005 | 0.481 | 0.422 | 0.043 | 0.577 | 0.203 |

| | | | | | | |
|---|---|---|---|---|---|---|
| F6P | 0.042 | 0.521 | 0.198 | 0.139 | 0.053 | 0.755 |
| FBP | 0.366 | 0.241 | 0.656 | 0.045 | 0.004 | 0.050 |
| DHAP | 0.109 | 0.168 | 0.327 | 0.156 | 0.010 | 0.002 |
| GAP | 0.408 | 0.032 | 0.017 | 0.094 | 0.205 | 0.367 |
| 3PG+2PG | 0.407 | 0.672 | 0.584 | 0.029 | 0.222 | 0.121 |
| PEP | 0.924 | 0.655 | 0.177 | 0.336 | 0.114 | 0.387 |
| Pyr | 0.549 | 0.085 | 0.525 | 0.346 | 0.041 | 0.031 |
| Lactate | nd | nd | nd | 0.310 | 0.047 | 0.375 |
| Citrate | 0.002 | 0.038 | 0.068 | 0.146 | 0.000 | 0.014 |
| 2KG | 0.846 | 0.091 | 0.672 | 0.294 | 0.008 | 0.021 |
| Succinate | 0.093 | 0.007 | 0.508 | 0.004 | 0.001 | 0.037 |
| Fumarate | 0.307 | 0.185 | 0.009 | 0.016 | 0.014 | 0.028 |
| Malate | 0.934 | 0.032 | 0.010 | 0.106 | 0.013 | 0.017 |
| 6PG | 0.003 | 0.647 | 0.896 | 0.090 | 0.002 | 0.023 |
| R5P | 0.187 | 0.082 | 0.385 | 0.565 | 0.099 | 0.962 |
| Ru5P | 0.270 | 0.703 | 0.062 | 0.337 | 0.316 | 0.141 |
| S7P | 0.005 | 0.070 | 0.068 | 0.925 | 0.147 | 0.116 |
| ATP | 0.256 | 0.332 | 0.073 | 0.139 | 0.186 | 0.457 |
| ADP | 0.767 | 0.603 | 0.831 | 0.043 | 0.169 | 0.919 |
| AMP | 0.104 | 0.411 | 0.124 | 0.002 | 0.026 | 0.032 |
| NAD | 0.779 | 0.863 | 0.006 | 0.000 | 0.096 | 0.039 |
| NADH | 0.476 | 0.396 | 0.042 | 0.117 | 0.001 | 0.251 |
| ATP/ADP | 0.299 | 0.158 | 0.298 | 0.006 | 0.128 | 0.494 |
| ATP/AMP | 0.012 | 0.125 | 0.049 | 0.013 | 0.121 | 0.013 |
| NADH/NAD | 0.000 | 0.407 | 0.101 | 0.031 | 0.000 | 0.125 |

**Table S4** *Saccharomyces cerevisiae* strains and plasmids used in this study

| Strains | Genotype | Sources |
|---|---|---|
| S288C (BY27002) | *MATα mal SUC2* | National BioResource Project (NBRP) |
| BY18615 | *MATα, ura3* | NBRP |
| YAS001 | BY18615[pGK426] | |
| YAS002 | BY18615[pAS1] | |
| YAS003 | BY18615[pAS2] | |
| **Plasmids** | **Description** | **Ref** |
| pGK426 | Yeast multi-copy type single-gene expression vector containing PGK1 promoter, PGK1 terminator, 2μ origin, and *URA3* marker | [17] |
| pGK426-QUEEN-2m | pGK426, expression of codon-optimized QUEEN-2m by PGK1 promoter | This study |
| pAS1 | pGK426, an open reading frame for expression of codon-optimized QUEEN-2m with the mitochondrial targeting signal derived from *S. cerevisiae* Cox4 at *N*-terminus by PGK1 promoter | This study |
| pAS2 | pGK426, two open reading frames for expression of codon-optimized QUEEN-2m with the mitochondrial targeting signal derived from *S. cerevisiae* Cox4 at *N*-terminus by PGK1 promoter | This study |

**Table S5** Primers for qPCR

| Genes | Primer sequences (5' to 3') (F: Forward and R: Reverse) |
|---|---|
| For *S. cerevisiae*, S288C | |
| *ACT1* | F：ACATCGTTATGTCCTTGGT<br>R：CCACCAATCCAGACGGAGTA |
| *HXT2* | F：CTTCGCATCCACTTTCGTG<br>R：AATCATGACGTTACCGGCAGCC |
| *HXT4* | F：ATGGAGAGTTCCATTAGGTCTAGG<br>R：ATAACAGCTGGATCGTCTGCGC |
| *PDC1* | F：CTTACGCCGCTGATGGTTA<br>R：GGCAATACCGTTCAAAGCAG |
| *CIT2* | F：CCTCCAGTTGGCTTATCGTG<br>R：CTGAAACGAATACCGTCTTCTG |
| *CAT2* | F：CCGTGGTTAAATTCATCGAAGC<br>R：ACTAAAGAAGACGGATCTGTGG |
| *FBP26* | F：GGAGATGTGCTACCGATATCC<br>R：GCCCACACGGCATACTTTTC |
| For human breast cancer cell, MCF7 | |
| *GAPDH* | F：ATGGAAATCCCATCACCATCTT<br>R：CGCCCCACTTGATTTTGG |
| *GLUT1* | F：AGGTGATCGAGGAGTTCTAC<br>R：TCAAAGGACTTGCCCAGTTT |
| *PFKP* | F：GGGCCAAGGTGTACTTCATC<br>R：TGGAGACACTCTCCCAGTCG |
| *LDHA* | F：GGACTTGGCAGATGAACTTG<br>R：TCAGAGAGACACCAGCAACA |
| *CPT1* | F：CCTCCAGTTGGCTTATCGTG<br>R：TTCTTCGTCTGGCTGGACAT |
| *CPT2* | F：GCAGATGATGGTTGAGTGCTCC<br>R：AGATGCCGCAGAGCAAACAAGTG |
| *PFKB4* | F：GGGTGCCTCTTGGCCTTAAA<br>R：GCCCACACGGCATACTTTTC |

**Table S6** Primers to construct pAS1 and pAS2

| Genes | Primer sequences |
|---|---|
| Queen-2m_Cox4N_invF | CAGCCACAAGAACTTTGTGTAGCTCTAG ATATCTGCTTATGAAGACCGTTAAGGTT AACATCACTAC |
| Queen-2m_Cox4N_invR | GCTTGAAAAATCTTATAGATTGACGTAG TGAAAGCATGTCGACGCTAGCGTTTTAT ATTTGTTG |
| AS1 fusion-F1 | GCGGTAATACGGTTATCCACAGAATC |
| AS1 fusion-R1 | CTTTGAGTGAGCTGATACCGCTC |
| AS1 QUEEN-F1 | TCAGCTCACTCAAAGAAAGATGCCGATT TGGGCG |
| AS1 QUEEN-R1 | TAACCGTATTACCGCAACGCAGAATTTT CGAGTTATTAAACTTAAAATACGC |
